# Oral cannabidiol administration in mice during pregnancy and lactation affects early postnatal body weight, fasting glucose, ingestive behavior, anxiety- and obsessive compulsive-like behaviors, and long-term object-memory in adult offspring in a sex-dependent manner

**DOI:** 10.1101/2024.07.10.602955

**Authors:** Martina Krakora Compagno, Claudia Rose Silver, Alexis Cox-Holmes, Kari B. Basso, Caroline Bishop, Amber Michal Bernstein, Aidan Carley, Joshua Cazorla, Jenna Claydon, Ashleigh Crane, Chloe Crespi, Emma Curley, Tyla Dolezel, Ezabelle Franck, Katie Heiden, Carley Marie Huffstetler, Ashley M. Loeven, Camilla Ann May, Nicholas Maykut, Alejandro Narvarez, Franklin A. Pacheco, Olivia Turner, Debra Ann Fadool

## Abstract

**Rationale:** The consequences of perinatal cannabidiol (CBD) exposure are severely understudied, but are important, given its widespread use and believed safety as a natural supplement.

**Objective:** The objective of this study was to test the health, metabolic, and behavioral consequences of perinatal CBD exposure on dams and their offspring raised to adult.

**Methods:** Primiparous female C57BL/6J mice were orally administered 100 mg/kg CBD in strawberry jam to expose offspring during gestation, lactation, or both using a cross-fostering design. Adult offspring were metabolically profiled using indirect calorimetry and intraperitoneal glucose tolerance testing. Adults were behaviorally phenotyped, video recorded, and mouse position tracked using DeepLabCut.

**Results:** CBD was detected in maternal plasma using LC-MS 10-min post consumption (34.2 ± 1.7 ng/ul) and peaked within 30 min (371.0 ± 34.0 ng/ul). Fetal exposure to CBD significantly decreased survival of the pups, and decreased male postnatal development, but did not alter litter size, maternal body weight or pup birth weight. We observed many sex-dependent effects of perinatal CBD exposure. Exposure to CBD during gestation and lactation increased meal size, caloric intake, and respiratory exchange ratio for adult male offspring, while exposure during lactation decreased fasting glucose, but had no effect on clearance. Adult female offspring exposed to CBD during lactation showed increased drink size. Perinatal CBD exposure increased obsessive compulsive- and decreased anxiety-like behaviors (marble burying, light-dark box, elevated-plus maze) in female mice, decreased long-term object memory in male mice, and had no effect on attention tasks for either sex.

**Conclusions:** We conclude that orally-administered CBD during pregnancy affects behavior and metabolism in a sex-dependent manner, and mice are differentially sensitive to exposure during gestation vs. lactation, or both. Because long-term changes are observed following perinatal exposure to the drug, and exposure significantly decreases survival to weaning, more research during development is warranted.

**Graphical Abstract:** 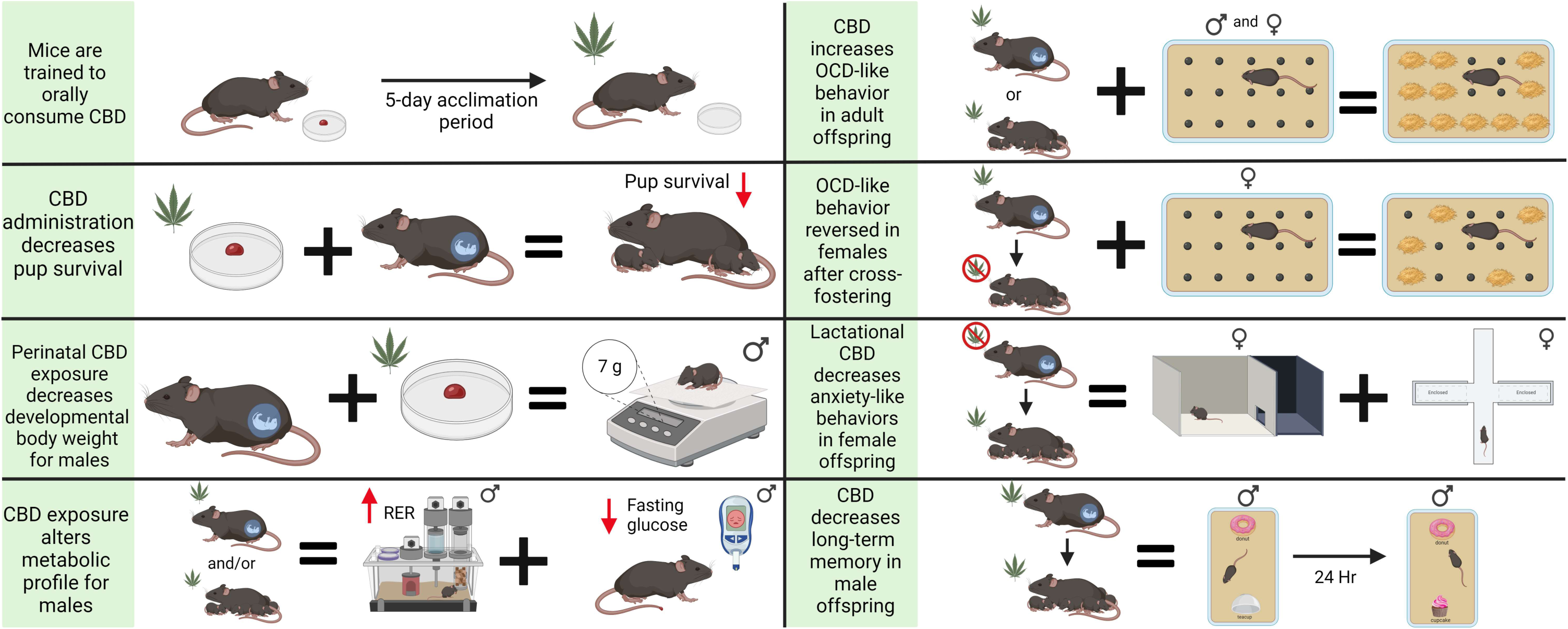

**HIGHLIGHTS:**

- Mice can be trained to orally consume CBD using strawberry jam as the vehicle.
- CBD administration to pregnant dams decreases pup survival to weaning age without significantly affecting maternal behavior.
- Perinatal CBD exposure decreases developmental body weight in males.
- Gestational or lactational CBD increases the respiratory exchange ratio (RER), increases mean meal and drink size, and reduces fasting glucose in a sex-dependent manner.
- CBD increases obsessive-compulsive like behavior in adult offspring, which could be eliminated in females by cross-fostering to a drug-free dam.
- Perinatal CBD selectively decreases anxiety-like behavior in females and decreases long-term object memory in males.

## 1. Introduction

The hemp plant, *Cannabis sativa,* contains hundreds of phytocannabinoids (multi-ring phenolic compounds derived from the cannabis plant) that may modulate activity in the central nervous system by interacting with the endocannabinoid system (Brown and Graves, 2013; Cristino et al., 2020; Schilling et al., 2020). The two most well-known phytocannabinoids are Δ9-tetrahydrocannabinol (Δ9-THC), the psychoactive component of marijuana, and cannabidiol (CBD), the major non-psychoactive and nonaddictive component of marijuana (Mackie, 2008; Calapai et al., 2020; Cristino et al., 2020).Because CBD does not induce the psychotropic effects clinically associated with Δ9 THC, it is publicly perceived as safe and free of harmful side effects (Grof, 2018; Bloomfield et al., 2019; American Society of Anesthesiologists, 2019). However, there is a growing demand for medical opinion on the use of *Cannabis* products based upon carefully designed research (Hill and Palastro, 2017; Bhamra et al., 2021). Simultaneously there is clinical interest in CBD due to its potential benefit as a natural antipsychotic, anti-nociceptive, anticonvulsant, antiemetic, anxiolytic, anti-inflammatory, antioxidant, and neuroprotective agent (Machado Bergamaschi et al., 2011; Hill and Palastro, 2017). Despite the usage of CBD to treat an array of ailments including anxiety, attention deficit disorders, ingestive disorders, neurodegenerative diseases, inflammation, pain, and nausea (Legare et al., 2022), there is not enough research attesting to the efficacy or safety of the drug. In fact, the only FDA-approved use of CBD is to treat specific forms of childhood epilepsy (Devinsky et al., 2017; Klotz et al., 2018; Ali et al., 2019). Although there are thought to be over 70 targets for cannabinoids (Ibeas Bih et al., 2015) the underlying cellular mechanisms of CBD have relied heavily on analogs or antagonists of the CB1/CB2 receptor pathways (Svizenska et al., 2008; Turner et al., 2017); leaving any potential long-term side effects of CBD and its metabolites largely understudied (Peng et al., 2022).

Given the rise of CBD use during pregnancy, where the drug is sought to mitigate pregnancy-related symptoms including nausea, insomnia, anxiety, and chronic pain (Volkow et al., 2012; Casagrande et al., 2015; Abuhasira et al., 2018; Sarrafpour et al., 2020), it is concerning that there are few studies exploring changes in brain development, cognitive function, altered behavior in offspring, or growth impairment upon gestational CBD exposure. Research investigating gestational CBD consumption is also difficult due to polysubstance use among participants (Gunn et al., 2016). In one clinical study, the authors surveyed use of *Cannabis* products by patients during pregnancy as 7%, whereas sampled uterine and cord blood demonstrated THC metabolites in 22% of these patients (Metz et al., 2019; Bidwell et al., 2020). Although it is known that cannabinoids cross the placenta as well as the blood-brain barrier, and have been detected in breast milk (Perez-Reyes et al. 1982; Bar-Oz et al. 2003; Shou et al. 2009; Ochiai et al., 2021), few preclinical studies have been performed (Wanner et al., 2021; Iezzi et al., 2022; Maciel et al., 2022; Swenson et al., 2023) that could advise safety precautions in the use of CBD during pregnancy. Studies have shown detrimental effects of prenatal *Cannabis* use – including early and long-lasting effects on cognitive function, placental abnormalities, fetal growth restrictions, and low birth weight and preterm delivery (Gunn et al., 2016; Natale et al., 2020). Thus, there is a need to study CBD use during pregnancy given its widespread public view of being “safe” (Grof, 2018; Bloomfield et al., 2019; American Society of Anesthesiologists 2019), while the FDA cautions against its use during pregnancy given the observed developmental effects in such *Cannabis* studies (Gunn et al., 2016).

Herein, we have designed a preclinical study in C57BL6 mice to examine the perinatal effects of CBD on both maternal health, and short- and long-term outcomes on offspring. We have used a cross-fostering design to separate any potential changes in maternal behavior from that of the physiological effects of the drug and evaluated early postnatal pup-dam interactions. This design also allowed us to examine differential effects of CBD exposure during gestation (embryonic development) vs. that of lactation (early postnatal development), or both. Because CBD is commonly consumed as an edible (Zhao et al., 2023), we developed a novel route of trained oral administration in mice using strawberry jam as a vehicle (Teixeira-Santos et al., 2021), which was also a stress-free route of administration during pregnancy. Offspring were developmentally monitored (survival and body weight), and as adults, they were metabolically profiled (total energy expenditure, locomotor activity, ingestive behavior, glucose clearance) and subjected to a battery of six behavioral tasks (buried marble, light dark box, elevated plus maze, short-term object memory, long-term object memory, and object-based attention tasks) designed to examine obsessive compulsive-like, anxiety-like, attention-deficit-like behaviors, and object memory in order to understand how fetal CBD exposure affects development and persists to changes in metabolism and behavior as an adult. Our study also utilized synthetic-derived CBD to confirm activity, known purity, and stability; and measured outcomes in both male and female offspring to compare any sexually dimorphic effects of perinatal administration of the drug. The importance of our reported outcomes is to help design future clinical trials that could advise safety precautions in the use of CBD during pregnancy, especially as there are no warning labels on CBD edible products (Russo, 2016).

## 2. Methods

### 2.1. Subjects

Drug administration was performed on 3-month-old primiparous female mice with C57BL6/J as the principal strain or strain background. A total of 68 drug- or vehicle-treated dams were crossed with male breeders to generate 49 litters and a total of 307 offspring. Of the offspring that were raised to adults to be phenotyped, 95 mice were analyzed for anxiety/memory-like behaviors, 115 were analyzed for metabolic profiles, and 95 were challenged for glucose clearance. Mice were either wildtype (Jackson Laboratories, stock number 000664, https://www.jax.org/strain/000664, RRID: IMSR_JAX:000664) or wildtype with an insertion of a YFP reporter previously shown to visualize subsets of neurons in the CNS without functional affect (Feng et al., 2000). All experiments in this study were approved, under protocol number #202300036, by the Florida State University (FSU) Institutional Animal Care and Use Committee (IACUC). These experiments were done in accordance with the guidelines set by the National Institutes of Health (NIH), the American Veterinary Medicine Association (AVMA), and the ARRIVE guidelines (du Sert et al., 2020). Upon weaning, all mice were singly housed with two sources of enrichment (house and bedding square) using open-style conventional cages. Breeding mice were kept on a regular 12-hr/12-hr light/dark cycle with lights on at 6:00 A.M. and lights off at 6:00 P.M. Once offspring were weaned, they were transitioned to a reverse 12-hr/12-hr light/dark cycle with lights off at 8:00 A.M. and lights on at 8:00 P.M. to facilitate behavioral testing in the dark cycle. All mice were given a standard diet (LabDiet 5001 Rodent Chow; 13.5% kcal from fat, https://www.labdiet.com/Products/StandardDiets/Rodents/index.html), and had access to food and water *ad libitum* in their home cages. For the duration of the behavioral experiments, mice were temporarily not given access to food or water for up to 2 hours (h).

### 2.2 Drug and Solutions

Cannabidiol (CBD) was obtained as synthetically derived from Purisys, LLC (Athens, GA; Batch NQS1951; NDC# 516342155) as purchased from and analytically certified by Emerald Scientific using mass spectrometry and infrared spectrometry. All CBD was of Pharmaceutical Grade, GMP Certified, with 100% purity factor rating, and without detection of Δ9-tetrahydrocannabinol (THC). CBD was orally delivered by mixing the drug in strawberry jam (Bonne Maman Strawberry Preserves, Publix Grocery Store, Tallahassee FL) at a final concentration of 100 mg/kg. The jam was administered as formed droplets ranging from 50 to 150 mg depending upon weight of the subject, and droplets were placed in the center of 35 mm corning dishes (Corning Inc., #430165, Corning, NY) (**Figure 2a)**. Drinking water was removed to introduce the dish to the front right corner of the home cage opposite the food remaining on the cage lid. Twenty to thirty milligrams of CBD was initially reconstituted in 100 ul of 100% ethanol prior to being diluted to a working concentration in the jam. The vehicle control solution contained 100 ul of 100% ethanol delivered at an equivalent jam volume as used for the drug (typically ranging from 850 to 950 mg jam). The ethyl alcohol was purchased from Pharmco-Aaper (Shelbyville, KY; Cat E200).

### 2.3 Jam Acclimation

One-hundred milligrams of strawberry jam was formed into droplets in 35 mm corning dishes as described above (**Figure 2a)**. Drinking water was removed from the cage and dishes were introduced to the right front corner of the cage floor opposite the food, which was retained on the cage lid. The time it took to complete consumption of the jam was recorded. Dishes were removed after consumption was complete or at 1 hour. Sixty minutes was the default recorded time for animals that did not consume all their jam within this cut off time. Dishes were presented daily for 5 days between 11:00AM and 2:00 PM, approximately 5 to 8 hours following the light cycle change.

### 2.4 Mass-Spectrometry / Liquid Chromatography of CBD in Maternal Plasma

Maternal plasma was prepared from whole blood and subjected to mass-spectrometry to quantify the kinetics of circulating CBD in the dams following oral administration. Following CBD consumption, mice were anesthetized with inhalation isoflurane and decapitated to collect trunk blood in chilled Eppendorf tubes that were precoated with 0.5M EDTA. An additional 2 ul of 0.5M EDTA/100 ul whole blood was added to the sample prior to centrifugation at 3,000 RPM in a 5425 Eppendorf (Eppendorf North America, Enfield, Massachusetts) for 15 minutes (min) without use of acceleration or brake. The plasma supernatant was stored at -80°C until analytical separation.

Samples were subsequently thawed in the dark and 100 μl was transferred to a new Eppendorf tube. Protein extraction was performed by adding 300 μl of cold acetonitrile to each sample, quickly vortexing, and then placing samples at -20°C for 3 hours (h). Samples were centrifuged at 12,000 g for 10 min and supernatant was transferred to a new tube for drying via speedvac. Following drying, samples were reconstituted in 50 μl of acetonitrile by alternating between sonicating on ice and vortexing for 2 h. Samples were covered with foil whenever possible to prevent light degradation. A 150 ng/ml spiked plasma served as an internal control and followed the same procedure as the samples. All standard curve samples (10, 25, 100, 300, and 500 ng/ml) were analyzed 3 times both before and after the sample run, and peak areas were averaged. All samples were analyzed in triplicate, and peak areas were averaged.

Concentration of CBD was analyzed in standard curve samples and in plasma samples using a high precision linear ion trap mass spectrometer (LTQ XL ThermoFinnigan, Atlanta, GA) in conjunction with liquid chromatography. Separation was optimized using a ThermoScientific Hypersil GOLD aQ (150 x 2.1 mm, 3 μm) column (column temperature: 40°C) and a gradient mobile phase containing 0.1% formic acid in water and 0.1% formic acid in acetonitrile. The electrospray ionization (ESI) was analyzed in positive mode using m/z range of 270-500, a spray voltage of 4 kV, and capillary temperature of 275°C. Samples were injected using a Thermo UltiMate 3000 series degasser and binary pump with 5 μl injection volume and separations maintained a flow rate of 0.300 μm/min (ThermoFisher). Chromatograms were analyzed using Xcalibur, v.2.2 SP1.48 software (ThermoFisher).

### 2.5 Cross Fostering and Health Monitored Metrics

To separate measurable effects of *in utero* drug exposure from that evoked by any changed maternal behaviors, it was necessary to cross foster mice. Dams were therefore bred in synchronized cohorts of 6 to 8 to increase the likelihood of paired birth. Cross fostering occurred between 24 and 48 hours of birth, across aged-matched or synchronized litters. Fostered pups would be removed from a pair of cages using individual clean gloves to prevent any transfer of odor or bedding material from the donor mother or her cage to the recipient mother’s cage. Pups were held at body temperature using a heating pad under the holding box. Urine-soaked litter and a few fecal pellets of the recipient mother’s cage was rubbed on the new pups to be introduced, prior to clustering all new pups in her nesting area at once.

### 2.6. Experimental Design

The experimental timeline is found in **Figure 1**. The female dams were acclimating to consuming jam from the dish for 5 days (week 1), after which they received drug or vehicle in the jam for a total of 14 days (weeks 2 – 3). Dams were then bred to males, and over the subsequent period (weeks 4 – 6), the dams continued to receive drug or vehicle over the course of gestation. Births were carefully monitored to accurately report number of births and the health status of early postnatal pups. Cross fostering occurred within the early window of birth (week 7), followed by continued oral administration throughout lactation (weeks 7 – 9). Pup retrieval assays occurred on three consecutive postnatal days 5 – 7 (week 8). Mice were weaned at postnatal day 23 (week 10) and then reared treatment free until 3-4 months of age (week 20). At this time, adult offspring first underwent metabolic assessment using indirect calorimetry (weeks 20-21), followed by intraperitoneal glucose tolerance testing (IPGTT) (week 22), and final behavioral testing (week 23 or 24). Health of the dams and the pups were monitored through this duration in terms of body weight, milk ribbons, and nesting/maternal behaviors.

**Fig. 1.**
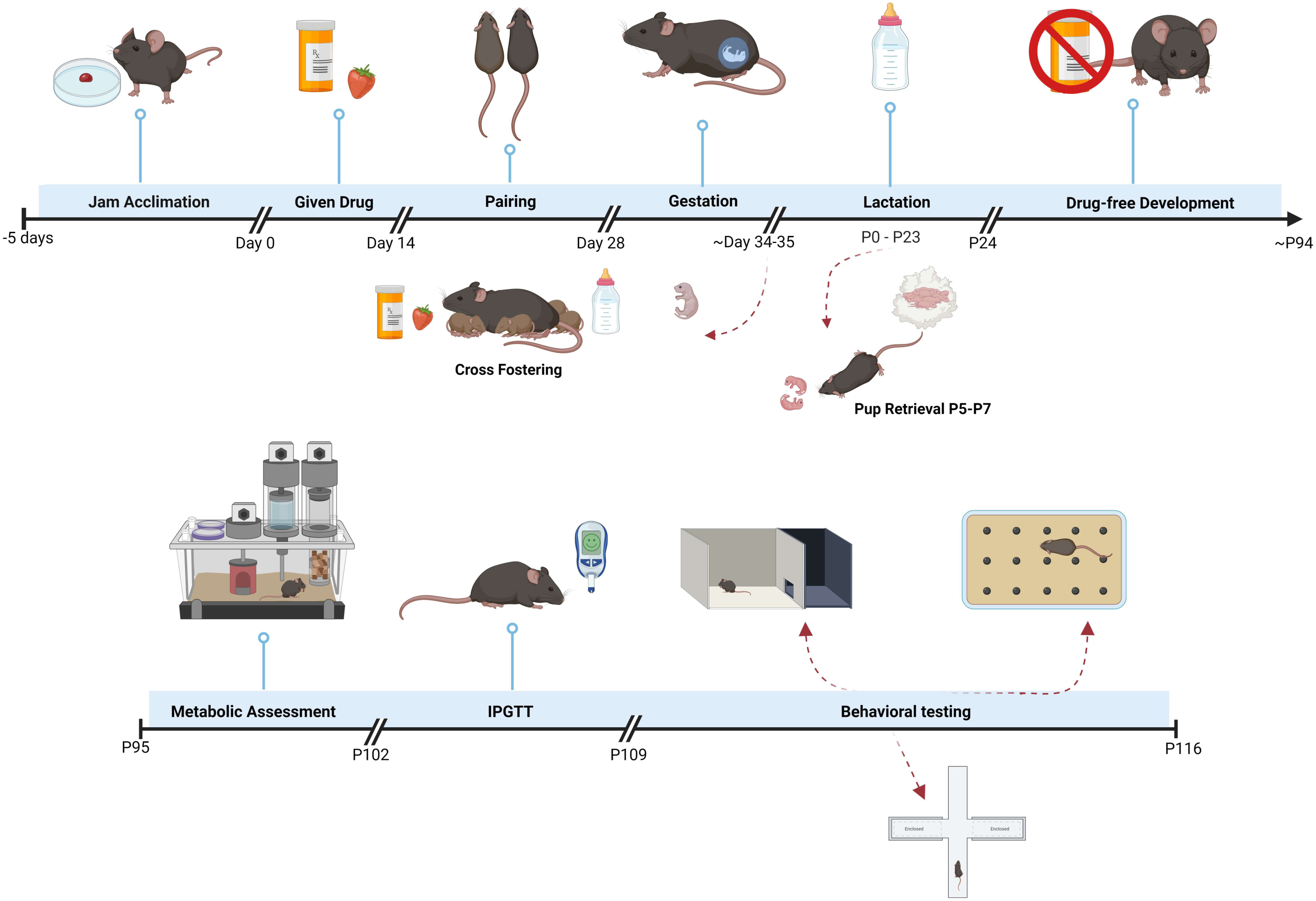
Experimental timeline for perinatal cannabidiol (CBD) exposure and testing of offspring in mice. Schematic demonstrating the timeline for dam drug exposure and behavioral and metabolic testing of offspring. Primiparous dams were acclimated to jam for 5 days (Jam Acclimation), given 100 mg/kg of CBD drug or solvent for two weeks (Given Drug), paired with a male for two weeks (Pairing), and given drug treatment up until weaning of pups (Gestation/Lactation). Offspring were raised drug free (Drug-free Development) until approximately postnatal day 94 when they underwent metabolic assessment (Metabolic Assessment) in a CLAMS system and glucose challenge (IPGTT) before behavioral phenotyping (Behavioral Testing). The light dark box (LDB) and elevated plus maze (EPM) evaluated anxiety-like behavior, the marble burying assay evaluated obsessive compulsive-like behavior, the object-based attention test evaluated attention-like deficit behavior (ADHD), and the object memory tests of 1- and 24-h duration evaluated any changes in short- or long-term memory. Behavioral tests were video recorded and manually scored by an observer that was blinded to the experimental condition. Video records were used to qualitatively map animal track movements using DeepLabCut software.

#### 2.6.1 Pup Retrieval

A pup retrieval assay was performed as per Winters et al, 2022 with minor modifications (Winters et al., 2022). Pups were withdrawn to a holding cage for 15 minutes that was outfitted with an underlining heating pad for room temperature maintenance. Pups were returned to the dam’s home cage as a cluster, and placed in the corner of the cage, diagonally opposite of the nest. Latency for first retrieval was scored as time until mother picked up first pup in her mouth. First retrieval was scored as time until the first pup was contained within the red igloo house surrounding the nest. Complete retrieval was scored as time until all pups were retrieved and contained within the red igloo house surrounding the nest. Complete retrieval was scored 60 minutes if dam had not completed retrieval of all pups within 1 hour, at which time, all pups were clustered back in the igloo for the dam. The order of which sex was retrieved was also noted in sequence by marking pups with a vertical (male) or horizonal bar (female) that was visible in the recording.

#### 2.6.2 Indirect Calorimetry

To assess modulation of whole-body metabolism due to perinatal CBD exposure, the mice were temporarily transferred to individual metabolic chambers at 3 months of age (**Fig. 1**). Mice were acclimated to metabolic chambers for 2 days prior to systems physiology measurement. Oxygen consumption (VO_2_; ml/kg/h), carbon dioxide production (VCO_2_; ml/kg/h), respiratory exchange ratio (RER), locomotor activity (beam breaks), food intake (kcal), and water consumption (ml) were tracked for ∼5 days using the Comprehensive Laboratory Animal Monitoring System (CLAMS^TM^; Columbus Instruments, Columbus, OH). Metabolic data were analyzed as the mean value over a continuous 72-hour interval acquired across the 5-day experimental window, sorted by light or dark cycle. During the period in the CLAMS^TM^, mice had *ad libitum* access to food and water. Food and water were delivered using overhead feeders attached to specialized electronic balances that monitored both disturbance and decrease in mass. A threshold of at least 10 seconds (s) of feeder disturbance and a minimum loss of 0.03 grams (g) of chow was required for a recorded meal bout. VO_2_ and VCO_2_ were normalized with respect to body weight in kilograms, calculated and averaged. RER was calculated as VCO_2_/VO_2_. Energy expenditure (EE) was calculated using the Lusk Graham equation (3.815 + 1.232 x RER) x VO2 (Lusk and Bois, 1924). Locomotor activity was continuously measured using optical beams along the X-axis of the cage (Columbus Instruments). Consecutive photo beam breaks were scored as ambulatory movement. Indirect calorimetry data were recorded in intervals using Oxymax software (Columbus Instruments). Each interval measurement represented the average value during a 30 s sampling period per cage.

#### 2.6.3 Intraperitoneal Glucose Tolerance Test

An intraperitoneal glucose tolerance test (IPGTT) was performed on the mice between 3-4 months of age (**Fig. 1**). Mice were fasted for 10 h starting at the beginning of the light phase, and then were injected with a volume of 25% glucose solution equivalent to 1 g of glucose per kg of body weight. A small incision was made on the tail, and blood samples were collected with a Contour Next Blood Glucose Monitoring System (Ascensia Diabetes Care US, Inc.; Parsippany, NJ) paired with Contour Next Blood Glucose Test Strips (Ascensia) to determine blood glucose levels at baseline (prior to injection) and at set timepoints 10, 20, 30, 60, 90, and 120 min following the injection. The area under the resulting curve was integrated (iAUC) per mouse and compared across perinatal treatment groups.

#### 2.6.4 Behavioral Tasks

Behavioral tasks were conducted to test anxiety-, attention-, and obsessive compulsive-like behaviors and object memory in mice. All mice received all six tests sequentially in the following order, namely 24-hr object memory recognition, light-dark box (LDB), elevated plus-maze (EPM), attention task, marble-burying, and then 1-hr object memory recognition. None of the anxiety-based behavioral tests we employed required hippocampal-dependent learning (Sartori et al., 2011), and any experience acquired in each task that might affect a subsequent task (novel object or attention tasks; (Antunes and Biala, 2012)) would have been equivalent for all mouse subjects due to the fixed order of the experiments. The sequential design also permitted us to reduce total number of mice in our study that otherwise would have been 6-fold.

Only one behavioral test was performed per day, except for day 2, when mice first completed their 24-hour memory test and were then introduced to the light-dark box, and day 3, when mice were first examined through the attention-task followed by direct introduction to the elevated plus maze. Dams had been randomly assigned to either drug or control vehicle, and their adult offspring now performed each of the six behavioral tests once and then were euthanized. Mice were phenotyped in cohort sizes of 8 to 16 mice, ensuring that all treatment groups and sexes were represented within a cohort. Treatment group sizes were matched through staged breeding as close as possible, but 13 mice had to be removed from the study due to failure in the Dixon Q test (n=4), failure in the object bias criteria (n=3), corrupt or incorrect video record (n=5), or one unexplained death prior to experimentation (n=1). During all animal handling and behavioral testing, investigators wore gloves, coats, masks, and protective shoe covers, and spoke minimally to avoid animal stress. Regardless of behavioral task, mice were acclimated to the new experimental chamber or testing room prior to data collection for varying durations as described individually below. All behavioral experiments were conducted during the dark cycle (08:00 - 20:00) in a room with temperature at 22°C and with humidity between 50 - 60%. All objects, boxes, and mazes were cleaned with 70% ethanol and air dried between subjects. All behaviors were filmed using a 16.6-megapixel Sony 4K Handycam with a 26.8 mm wide angle lens and still image recording (FDR-AX53; Best Buy, Tallahassee, FL) mounted to a Sony tripod using a 1K Gorillapod (Joby Aviation, Santa Cruz, CA). Films were uploaded to private access YouTube (YouTube, San Bruno, CA) so that researchers could score activity blinded to the drug or genotype condition. Following download from YouTube, QuickTime Player (Apple, Inc., Cupertino, CA) and Movies and TV (Microsoft, Redman, WA) were used as the platforms to manually score respective behavior activities by Mac and Windows users, respectively. Films were stored on Passport External Drives (Western Digital, San Jose, CA) and are available through request.

##### 2.6.4.1 Marble Burying Test

The marble burying test assesses anxiety-like and obsessive-compulsive-like behavior in mice (Nicolas et al., 2006; Dixit et al., 2020). We followed the procedure as previously used by Marks et al. 2009 and described by others (Marks et al., 2009; Dixit et al., 2020). Briefly, mice were taken from their home cage and placed in a standard rat cage/box (45 cm [L] × 23 cm [W] × 20 cm [H]) with 3 cm of bedding to acclimate for 30 minutes. The cage lid, water, and food were removed and replaced with a sheet of plexiglass (Amazon, Seattle, WA) to prevent distraction and allow video recording of behaviors. For testing, 18 black metallic marbles (Amazon) were placed in a 6-marble x 3-marble grid (**Figure 10a)**. To initiate a trial, mice were placed in the center of the test cage and were allowed to move freely for 30 minutes. A picture was taken of the cage before the marbles were touched and after the mouse was recovered to its home cage to ensure accuracy of the marbles-buried count. To be considered buried, the surface of the marble had to be covered ⅔ of the way with bedding. Following the trial, marbles were recovered from the bedding and washed with Versa-Clean (VWR, Radnor, PA) diluted in water, sprayed with 70% EtOH, and air dried before being used again.

##### 2.6.4.2. Light-dark box

The light-dark box (LDB) assesses anxiety-like behavior in rodents that avoid time spent in the illuminated side (Hascoët et al., 2001; Bourin and Hascoët, 2003; Takao and Miyakawa, 2006; Bourin, 2015). In addition to time spent in each compartment, the number of transitions between the two compartments can be scored as an index of general locomotor activity. Latency of first movement to the opposite compartment than initially placed is also an index of anxiety. The LDB testing chamber was a standard rat cage (45 cm [L] × 23 cm [W] × 20 cm [H]) painted black on one side and painted white on the other side. A black divider was positioned in the center with a small entry door (7 cm [L] x 7 cm [H]) to allow ease of mouse movement and transition between the created two compartments. A 60-watt light bulb was hung 70 cm directly over the center of the white chamber and a sheet of plexiglass again served as the cage lid to prevent distraction and allow video recording of behaviors. The LDB protocol was as previously performed (Marks et al., 2009; Huang et al., 2018a). Briefly, mice were acclimated to the testing room 30 minutes prior to the experiment. Mice were then transferred from their home cage into the LDB apparatus by placing the mouse in the light compartment. The trial duration was 5 minutes, where the time to first latency to the dark, number of total transitions, and time spent in each compartment were scored. Entrance into a compartment was defined as all four paws of the mouse crossing into that compartment. The LDB was cleaned using Kimwipes (VWR) and 70% ethanol between each mouse trial.

##### 2.6.4.3. Elevated plus maze

The elevated plus maze (EPM) assesses anxiety-like behavior in rodents due to their proclivity for movement to enclosed spaces and their unconditioned fear of heights (Montgomery, 1955; Pellow et al., 1985; Walf and Frye, 2007). The EPM can be used to evaluate potential changes in anxious behavior based upon changed distribution of time spent in the maze compartments (Komada et al., 2008). Our apparatus consisted of four arms (35 cm [L] x 5 cm [W] each) raised to a height [H] of 45 cm from the ground. Two arms were completely flat and lacked barriers (open arms) and two arms had 15 cm tall barriers (closed arms). The procedure was as performed previously (Huang et al., 2018a), whereby mice were acclimated to the experiment room for 30 minutes prior to experimentation. Mice were then introduced to the middle of the plus maze facing the open arm and allowed to freely explore the apparatus for a 5-minute duration. The time spent in the open and closed arms and the total number of transitions were scored. Entrance into an arm was defined as all four paws of the mouse crossing into that arm. The EPM was cleaned with Kimwipes and 70% ethanol between mouse subjects.

##### 2.6.4.4. Short- and long-term memory testing

Mice were tested for short- (1 h) and long-term (24 h) object recognition that are tests for memory in rodents as previously performed (Marks et al., 2009; Antunes and Biala, 2012; Tucker et al., 2012; Huang et al., 2018a). Briefly, mice were acclimated for 30 minutes in a standard rat cage (45 cm [L] × 23 cm [W] × 20 cm [H]) filled with a layer of bedding and with the cage lid and food/water removed and replaced with a sheet of plexiglass as described above. Following acclimation, two unfamiliar objects (object 1, object 2) were placed at opposite ends of the cage and mice were allowed to explore for a duration of 5 minutes. Then, either 1 h (short-term memory) or 24 h (long-term memory) later, one familiar object (object 1) and one novel object (object 3) were alternatively placed in the cage and the mice were given opportunity to explore for a second 5-minute duration. The mouse was considered to be exploring the object when all of the following criteria were met: the mouse was oriented toward the object, the nose was within 2 cm of the object, and both of these criteria were met for at least 1 second. Chewing or standing on an object did not count toward exploration time. A recognition index (RI) was calculated based upon time spent with the novel object, or RI = object 3/(object 1 + object 3), whereby a higher RI was associated with better memory retention. In order to assure equal time exploration of the initial object 1 and object 2, an object bias score was calculated, or OBS = object 1/(object 1 + object 2). Mice exhibiting an OBS below 0.20 or above 0.80 (i.e. signifying avoidance or attraction of object 1, respectively) were noted and excluded from the subsequent 1 h or 24 h object memory test. The objects used were plastic toys of similar size but different shapes (McDonald’s, Chicago, IL), and they were cleaned using Kimwipes and 70% ethanol between mouse subjects.

##### 2.6.4.5. Object-based attention testing

Attention deficit-like behavior (ADHD) in rodents can be assessed with an object-based attention test as originally designed (Alkam et al., 2011; Ishisaka et al., 2012) and as previously performed (Huang et al., 2018a). The object-based attention test is similar in design as the object memory paradigm, but it is of shorter duration and more objects are presented. The testing chamber is larger and has two unequal compartments separated with a removable divider: a larger chamber (40 cm [L] x 40 cm [W] x 22 cm [H]) and a smaller (20 cm [L] x 40 cm [W] x 22 cm [H])) chamber. The testing chamber was filled with a layer of bedding, and the cage lid and food/water were removed and replaced with plexiglass as described above. Mice were first acclimated to the full chamber (no divider) for 10 minutes. Mice were gently coaxed into the larger compartment, the divider was inserted, and mice were allowed to explore for 10 minutes. Mice were then coaxed into the smaller compartment and allowed to explore for an additional 10 minutes. Following this full acclimation period of 30 minutes, five wooden shapes (Walmart, Bentonville, AR) were placed in the larger compartment and mice were allowed to explore the objects for 3 minutes. Mice were then gently coaxed into the smaller compartment where they were presented with two objects, one was a novel object (NO) and the second was a familiar object (FO) matching one of the five shapes from the larger compartment, whereby mice were allowed to explore for 3 minutes. Criteria for object exploration were the same as described previously for the short- and long-term memory testing. RI was calculated as NO/(NO + FO), based upon time spent exploring the objects. The mouse was considered to be exploring the object when all of the following criteria were met: the mouse was oriented toward the object, the nose was within 2 cm of the object, and both of these criteria were met for at least 1 second. A lower RI indicated increased attention-like deficit behavior. In the object-based attention test, the box and shapes were cleaned with 70% ethanol and new bedding was added for each mouse subject.

### 2.7. Mouse traces

Videos from behavioral tasks were analyzed to examine differences in patterns of movement. Tracking information was determined using DeepLabCut (Mathis et al., 2018), whereby we manually marked five body parts (nose, neck, spine, hips, and tail base), in 20 representative frames per video, generating a training set for a convolutional neural network designed to locate these five body parts on a frame-by-frame basis. X and Y coordinates of DeepLabCut tracking data were imported into R studio (v 4.3.2) and traces for each behavior were generated using custom scripts. Custom scripts are available upon request.

### 2.8. Statistical analysis

Data manually scored from video playback were organized in Excel (Microsoft Office 365 Suite), then analyzed and graphed in Prism v9.0 (GraphPad Software, Inc.), and finally compiled to Photoshop CS4 (Adobe, San Jose, CA) to create figure layouts. Prior to performing any statistical comparisons, data were first analyzed with the Dixon’s Q test to identify any outliers. Data were then checked for normal distribution and homogeneity of variance using the F_max_ test. Analysis of data collected across the behavioral phenotyping experiments infrequently identified outliers (data from three mice in the LDB, and one mouse in the first latency test) nor did any collected data violate homogeneity of variance (fail the F_max_ test). Data collected in the CLAMS had infrequent outliers (three male mice light cycle, one male mouse dark cycle, 5 female mice light cycle, and 2 female mice dark cycle) distributed across the 13 metabolic parameters that were measured per light cycle and in each sex, and no values failed the F_max_ test. The iAUC computed for glucose clearance in the IPGTT was analyzed by comparing the integrated areas across perinatal treatment groups using a 1-way analysis of variance (1-w ANOVA) at the 95% confidence level (α ≤ 0.05). The Bonferroni method for multiple comparison testing was used as the *post-hoc* analysis in this, and subsequent applied ANOVAs, to make mean-wise comparisons between treatments. Pup retrieval data were analyzed using a mixed repeated measure (RM) 2-way analysis of variance (2-w ANOVA) with drug and postnatal age (RM) as factors, and drug x age interactions at the 95% confidence level (α ≤ 0.05). Maternal health outcome data were analyzed by Student’s *t*-test for two sample comparisons (drug vs. vehicle). Maternal body weight over gestational day was analyzed using a RM 2-w ANOVA with drug and gestation day (RM) as factors. A non-parametric equivalent to the 1-w ANOVA (Kruskal Wallace) was applied to percent death prior to weaning data because the metric was not normally distributed (percentage data). The developmental pup weights were analyzed with an ordinary 2-w ANOVA rather than a RM design, because the P1 pups could not be tagged for paired subsequent body weight determination (ear tagging began at P10). All CLAMS metabolic data were analyzed within sex and within light cycle to compare a given metric across the four drug treatment groups (1-w ANOVA).

The number of buried marbles, number of transitions and time to first latency in the LDB, and number of transitions in the EPM were compared across perinatal treatment groups using a 1-w ANOVA (α ≤ 0.05). Mouse behavior in the EPM and LDB was analyzed using a RM 2-w ANOVA with drug and location (RM) as factors, and drug x location interactions at the 95% confidence level (α ≤ 0.05). Recognition Indexes (RI) computed in the short- and long-term object memory and object-based attention tasks were compared across perinatal treatment groups using a Kruskal Wallis (non-parametric equivalent to the ANOVA) appropriate for percentage data. All data were analyzed within sex as independent cohorts. All reported values in the text and figures are mean ± standard deviation (SD). Sample sizes are reported as individual data points in the graphs and represent number of mice. Individual *F* statistic and/or *p* values are reported for each behavioral experiment within the text and the corresponding graph, as described in the results section. Additionally, a full summary of all applied statistics, sample size, and experimental question as sorted by figure number is reported in **Supplementary Table 1**. Applied statistics and *p* values for metabolic parameters generated from the CLAMS experiments are broken out separately as reported in **Supplementary Table 2** and described in the results section.

## 3. Results

### 3.1. Jam Acclimation and CBD Detection in Maternal Plasma

#### 3.1.1 Oral Administration of CBD in Mice

Most adult mice were able to learn to consume CBD contained in strawberry jam within the course of 5 days of acclimation. In a sample of 37 mice treated with oral administration, only 1 mouse failed to learn (3%). On average, mice took 40.6 ± 3.6 minutes to investigate and consume a ∼100 mg droplet of jam upon first presentation, but within 5 days, the mean time of consumption significantly shortened to 3.9 ± 0.7 min (**Figure 2A-B**; 1-w RM ANOVA, F(4, 144) = 46.93, *p* < 0.0001, with a Bonferroni’s post-hoc test, \*\*\*\**p* < 0.0001 for all time points compared with day 1). Furthermore, as mice transitioned from initial jam acclimation to daily jam consumption (**Figure 1**; solvent vs. CBD drug), most mice consumed the jam droplet in less than 15 seconds once approaching the dish. This demonstrated that oral drug administration using strawberry jam was an effective route of administration. Interestingly, in a separate cohort of younger mice (P24), animals failed to learn to eat jam, and even after 2 weeks of daily presentation, the time of consumption ranged from 40 min to the default score of 60 min for failure to consume (mixed 2-w ANOVA, sex and time as factors, no main effect of time, F(4.199, 29.40) = 1.818, *p* = 0.0659). Therefore, age of mice was an important factor for learning the oral administration, and only dams that were 3 months of age were subsequently used in our study.

**Fig. 2.**
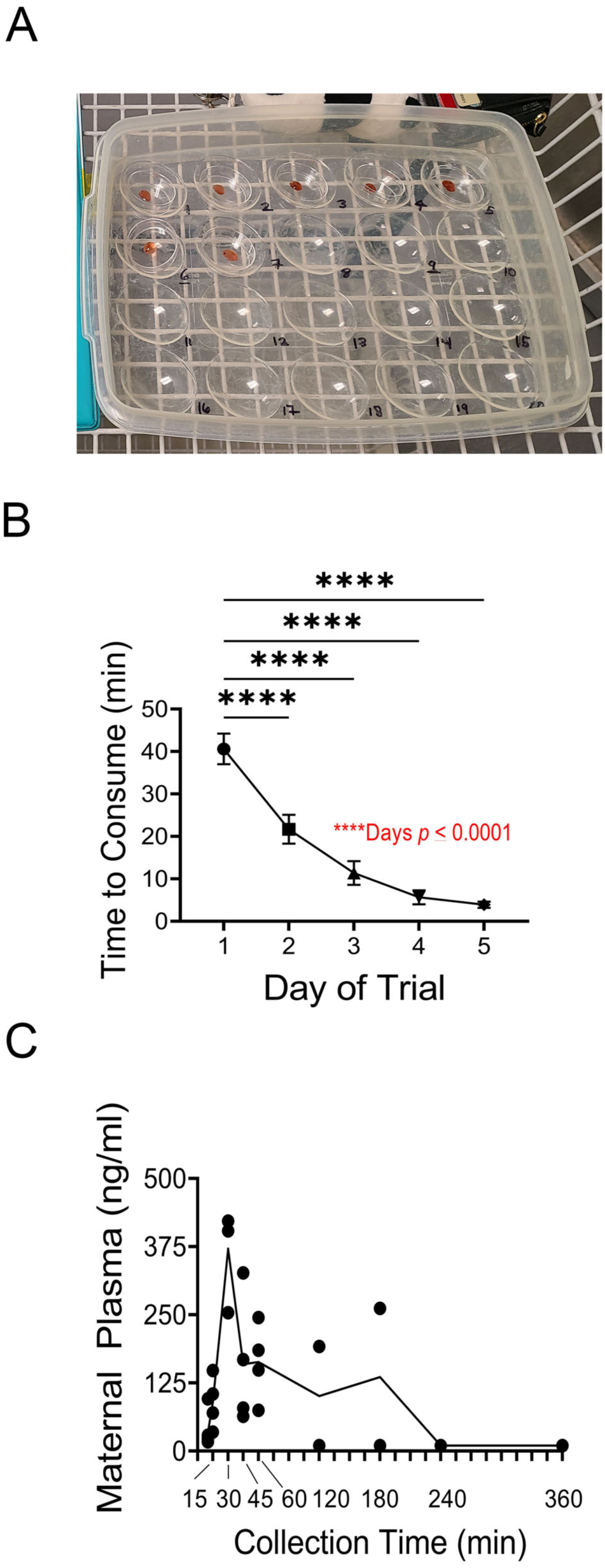
Mice can be trained to orally consume CBD in a dollop of strawberry jam. A, Representative image of 35 mm dishes with strawberry jam that were either mixed with CBD drug or ethanol solvent and placed in the mouse’ home cage for oral consumption. B, Line graph of time to consume jam following the course of 5 days of jam acclimation. Following 1 week of Jam acclimation, mice consumed the dollop of jam within 15 seconds. 1-way RM ANOVA, *Bonferroni’s* post-hoc \*\*\*\**p* < 0.0001 C, Line graph plotting concentration of CBD in maternal plasma over time since oral consumption. Data represent individual dams that orally consumed 100 mg/kg CBD in strawberry jam (time 0) and plasma concentration of CBD determined through collection of truck blood at designated time points.

#### 3.1.2. Pharmacokinetics Following Oral CBD Administration

A total of 24 retired dam breeders were used to harvest maternal plasma following 100 mg/kg administration of CBD. The concentration of CBD in the plasma was determined using mass-spectrometry / liquid chromatography measures – and was found to be surprisingly rapidly detected in the circulation following oral consumption. CBD could be detected as rapidly as 10 min post consumption (34.2 ± 1.7 ng/ml), it peaked within 30 min (371.0 ± 34.0 ng/ml) and was below the limit of detection levels in our assay by 4 hours (< 10 ng/ml) (**Figure 2C**).

### 3.2. Pup Retrieval – Maternal Behavior

Maternal behavior in response to nest disturbance (**Figure 3A**) was not notably different across the four treatment groups: 1 – Jam throughout gestation and lactation, 2 – CBD throughout gestation and lactation, 3 – Jam during gestation and CBD during lactation, or 4 – CBD during gestation and Jam during lactation. The range for first latency (see methods) by the dam was quick, but broad (1 to 47 s), regardless of the treatment group of the pups. There was a main effect of postnatal day, in that mothers became increasingly more rapid in subsequent, first latency retrievals with repeated trials, but there was no effect of treatment group (**Figure 3B**, 2-w mixed RM ANOVA, postnatal age F (1.692, 42.29) = 5.707, \*\**p* = 0.0091; treatment F (3, 26) = 0.5425, *p* = 0.6575). There was also no significant effect of treatment for the total time to collect an entire litter back to the nest (**Figure 3C**, 2-w mixed RM ANOVA, treatment F (3, 26) = 0.3891, *p* = 0.7618).

**Fig. 3.**
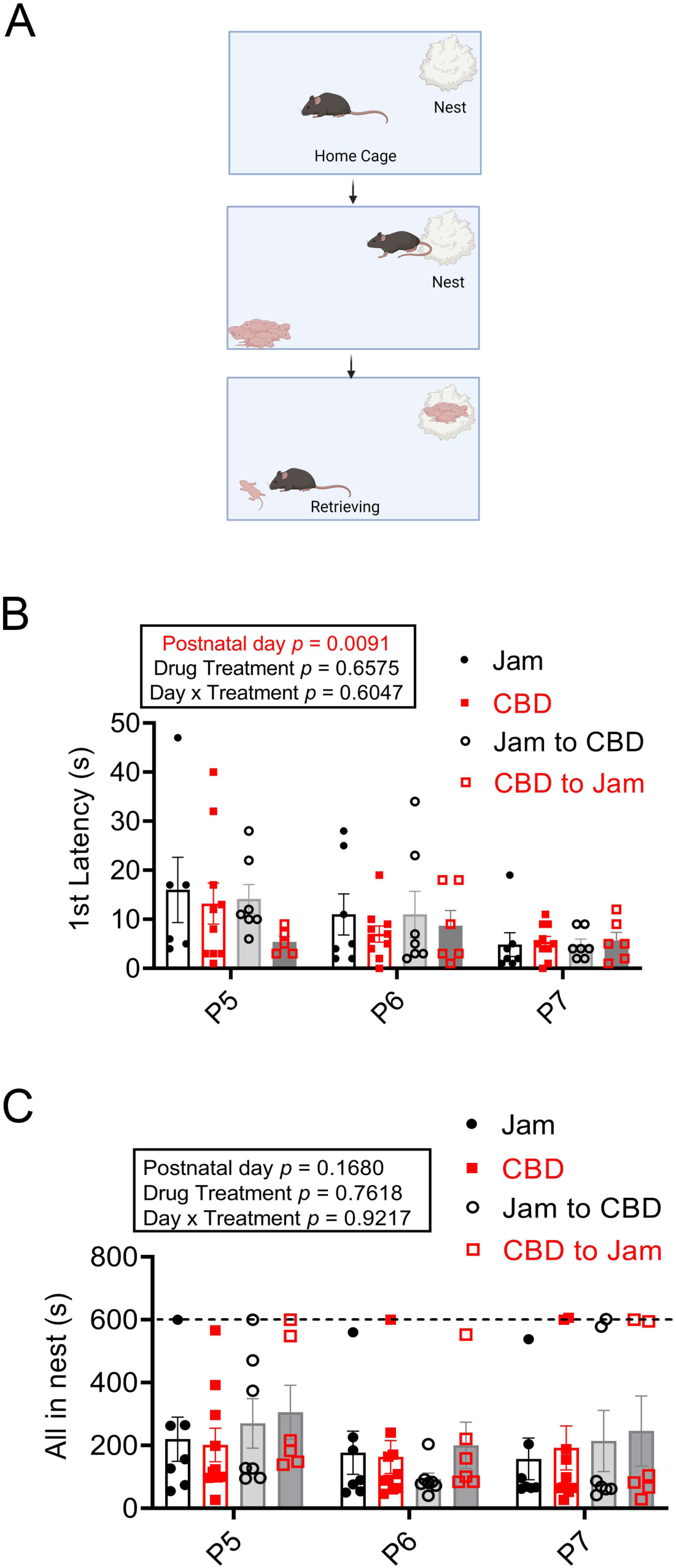
Perinatal CBD exposure does not significantly affect maternal behavior in a pup retrieval test. A, Schematic of pup retrieval test. (top panel) Dams are separated from pups for 15 min. (middle panel) Pups are reintroduced to the cage diagonally from the nest and (bottom panel) the time of the dam’s first contact (First Latency, B) and collection of all pups to the nest (All in Nest, C) are scored. Bar graph of B, First Latency and C, All in Nest plotted against postnatal age (P) of the pup in days. Data represent individual dams that were tested with a repeated measure over the three sequential days. B, 2-w mixed RM ANOVA, main effect of postnatal day \*\**p* < 0.01, no significant effect of drug treatment or day x treatment interaction. C, 2-w mixed RM ANOVA, no significant effect of day, drug, or day x treatment interaction. Legend here and in subsequent figures: Jam = (control) strawberry Jam plus ethanol solvent, CBD = 100 mg/kg CBD in strawberry Jam, Jam to CBD = cross-fostered to achieve administration of Jam during gestation and then CBD during lactation (after birth), CBD to Jam = cross-fostered to achieve CBD during gestation and then Jam during lactation (after birth).

### 3.3. Health and Metabolic Status Following Perinatal CBD Exposure

#### 3.3.1. Maternal Weight Gain, Litter Size, Length of Gestation, and Pup Survival

A total of 41 pregnant dams and their litters were monitored for health or metabolic status following perinatal CBD or Jam exposure. Dam maternal weight gain significantly increased with gestation day (**Figure 4A**, 2-w mixed RM ANOVA, *p* < 0.0001). There was no main effect of drug treatment (*p* = 0.1752), nor was the total maternal weight gain significantly different between solvent- and CBD-treated dams (calculated as body weight (BW) at gestational day 20 – initial BW upon crossing with a male; **Figure 4A, inset**; Student’s *t*-test, *p* = 0.7605; solvent = 12.5 ± 2.0 g vs. CBD = 12.3 ± 2.0 g). There was a modest drug x day interaction (**Figure 4A**, *p* = 0.0478) but without any significant difference in the *post-hoc* analysis (*p* ≥ 0.05). CBD did not affect the length of gestation (**Figure 4B**, Student’s *t*-test, *p* = 0.7553; solvent = 21.6 ± 0.3 days vs. CBD = 21.4 ± 0.5 days). While CBD did not affect the initial size of the litter (**Figure 4C**, Student’s *t*-test, *p* = 0.9635; solvent = 6.6 ± 1.6 pups vs. CBD = 7.5 ± 1.6 pups), gestational CBD exposure significantly affected survival of the offspring to weaning age (**Figure 5A**, Kruskal-Wallis, *p* = 0.0395). When exposed to CBD during gestation and lactation 37.9 ± 2.3 % of the litter did not survive to weaning age (P23). In contrast, when pups were exposed to Jam (solvent), 8.1 ± 2.3% of the litter did not survive (*p* = 0.0439, multiple comparison, *post-hoc* test). Early postnatal exposure to CBD during lactation (Jam to CBD treatment) had a survival rate that was not significantly different than Jam (solvent) treatment (*p* > 0.9999), and cross-fostering of drug-exposed pups *in utero* to drug-free dams during lactation (CBD to Jam treatment) also was not significantly different than Jam (solvent) treatment (*p* = 0.1431), indicating early embryonic exposure to drug was the most detrimental variable for pup survival rate.

**Fig. 4.**
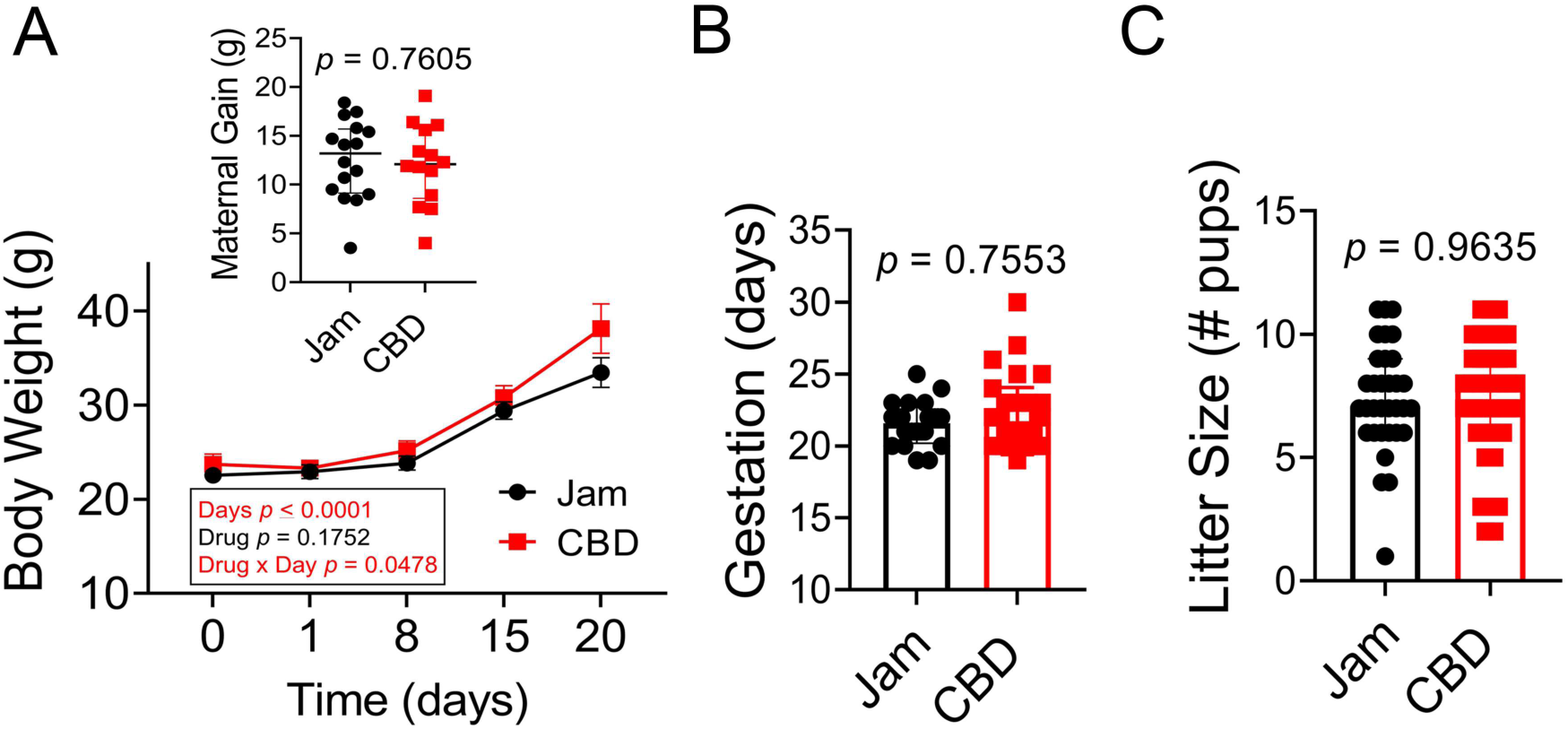
CBD administration did not affect total maternal weight gain, duration of gestation, or size of the litter. A, Line graph of the mean (± SEM) maternal weight gain over time. Dam maternal weight significantly increased with gestation day without a main effect of drug treatment. 2-w mixed RM ANOVA, ****p < 0.000; inset comparing change in maternal weight by day 20, data represent individual dams plotted in the main line graph. B, Bar graph of gestational length defined as days until birth following introduction of a male. C, Bar graph of litter size. A inset, B-C, Student’s *t*-test, \**p* < 0.05, data represent individual dams with mean (± SEM).

**Fig. 5.**
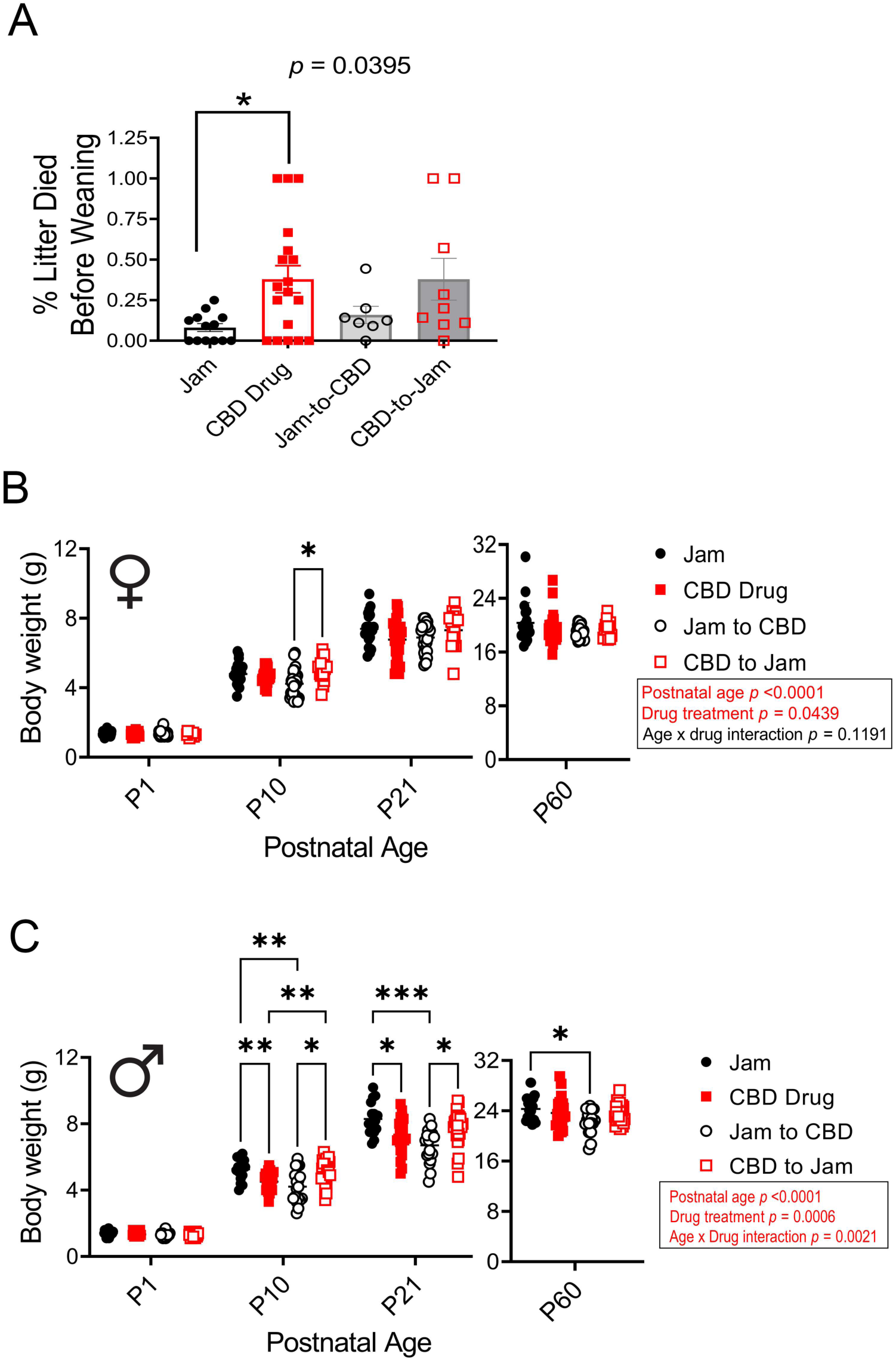
Perinatal CBD exposure decreases pup survival and affects early postnatal body weight. A, Bar graph of the percentage of the litter that died prior to P24 weaning age following one of the four drug treatments. Data represent individual litters (± SEM). Kruskal-Wallis, \**p* < 0.05. Scatter plot of the body weight for B, female and C, male offspring following one of the four drug treatments sorted by postnatal age. 2-way RM ANOVA, *Bonferroni* post-hoc \**p* < 0.05, \*\**p* < 0.01, \*\*\**p* < 0.001

#### 3.3.2. Developmental Growth and Glucose Clearance in Adult Offspring

Interestingly, in a comparison of pup BW in surviving animals exposed *in utero* to either Jam or CBD treatment, BW was not significantly different at postnatal day (P) 1 (females - solvent = 1.4 ± 0.03 g, CBD = 1.3 + 0.02 g, Jam to CBD = 1.4 ± 0.03 g, CBD to Jam = 1.3 ± 0.03 g; males – solvent = 1.4 ± 0.04 g, CBD = 1.3 ± 0.02, Jam to CBD = 1.3 ± 0.04 g, CBD to Jam = 1.3 ± 0.02 g), but BW was significantly different in drug-exposed pups by age P10 to P21 in a sex-dependent manner (**Figure 5B, C**). For the female mice, there was both an age and treatment effect during a short period at P10, where mice exposed to CBD during gestation weighed more than those exposed to CBD during lactation (**Figure 5B**; 2-w RM ANOVA, drug treatment F (3, 91) = 2.809, *p* <0.0439; postnatal age F (1.439, 131) = 5345, *p* <0.0001; Bonferroni post-hoc test, \**p* = 0.0128; P10 solvent = 4.8 ± 0.14 g, CBD = 4.7 ± 0.07 g, Jam to CBD = 4.2 ± 0.14 g, CBD to Jam = 5.0 ± 0.18 g; *p* > 0.05; P21 solvent = 7.4 ± 0.21 g, CBD = 6.8 ± 0.20 g, Jam to CBD = 6.9 ± 0.15 g, CBD to Jam = 7.3 ± 0.27 g). For the male mice, however, animals exhibited both an age and treatment effect throughout P10 to P21, that persisted for some treatment groups as adults (**Figure 5C**; 2-w RM ANOVA, drug treatment F (3, 78) = 6.440, *p* = 0.006; postnatal age F (1.565, 122.1) = 8504; age x drug interaction F (9, 234) = 3, *p* = 0.0021). For the male mice, the *post-hoc* test oppositely showed that perinatal CBD exposure resulted in a reduction in BW compared to that of Jam-treated mice (Bonferroni post-hoc, Jam vs. CBD during gestation and lactation, \*\**p* = 0.0040 at P10 and \**p* = 0.0100 at P21; Jam vs. CBD during lactation, \*\**p* = 0.0047 at P10, \*\*\**p* = 0.0002 at P21, and \**p* = 0.0267 at P60; test, \**p* = 0.0128; P10 solvent = 5.3 ± 0.16 g, CBD = 4.5 ± 0.12 g, Jam to CBD = 4.2 ± 0.23 g, CBD to Jam = 5.2 ± 0.14 g; P21 solvent = 8.3 ± 0.23 g, CBD = 7.2 ± 0.23 g, Jam to CBD = 6.7 ± 0.24 g, CBD to Jam = 7.7 ± 0.21 g; P60 solvent = 24.3 ± 0.26 g, CBD = 23.7 ± 0.51 g, Jam to CBD = 22.3 ± 0.45 g, CBD to Jam = 23.6 ± 0.30). Due to observed changes in BW with early postnatal age, we wondered if there would be a persistent change in an animal’s ability to clear a glucose challenge as an adult. Adult male mice, but not female, had a significantly lower fasting glucose (**Figure 6A-B**; 1-w ANOVA, females F(3, 41) = 1.928, *p* = 0.1401 and males F(3, 44) = 3.983, *p* = 0.0135; Bonferroni *post-hoc* test, \**p* = 0.0443 Jam vs. CBD during lactation, \**p* = 0.0127 CBD during lactation vs. CBD during gestation). Both sexes had a significant time x treatment interaction (2-w RM ANOVA, female F(18, 247) = 3.959, *p* ≤ 0.0001, male F(18, 269) = 1.657; *p* = 0.0470) in the time course of their ability to clear a glucose challenge (**Figure 6C, 6E respectively**). Female adult mice that were exposed to CBD *in utero* had a higher elevation of glucose at 20 minutes but then cleared the glucose challenge more quickly between 90 and 120 minutes compared with that of male mice that showed no difference in the timing of glucose clearance, and only changes in their fasted levels at time -10 minutes (**Figure 6E, 6F**). Albeit some kinetic differences in glucose clearance in the females, this did not result in an overall change in clearance ability, as reflected in a lack of significant difference for either sex for computed integrated area under the curve data (iAUC, **Figure 6D, 6F**; 1-w ANOVA, females F(3, 34) = 1.582, *p* = 0.2118 and males F(3, 45) = 2.205, *p* = 0.1005).

**Fig. 6.**
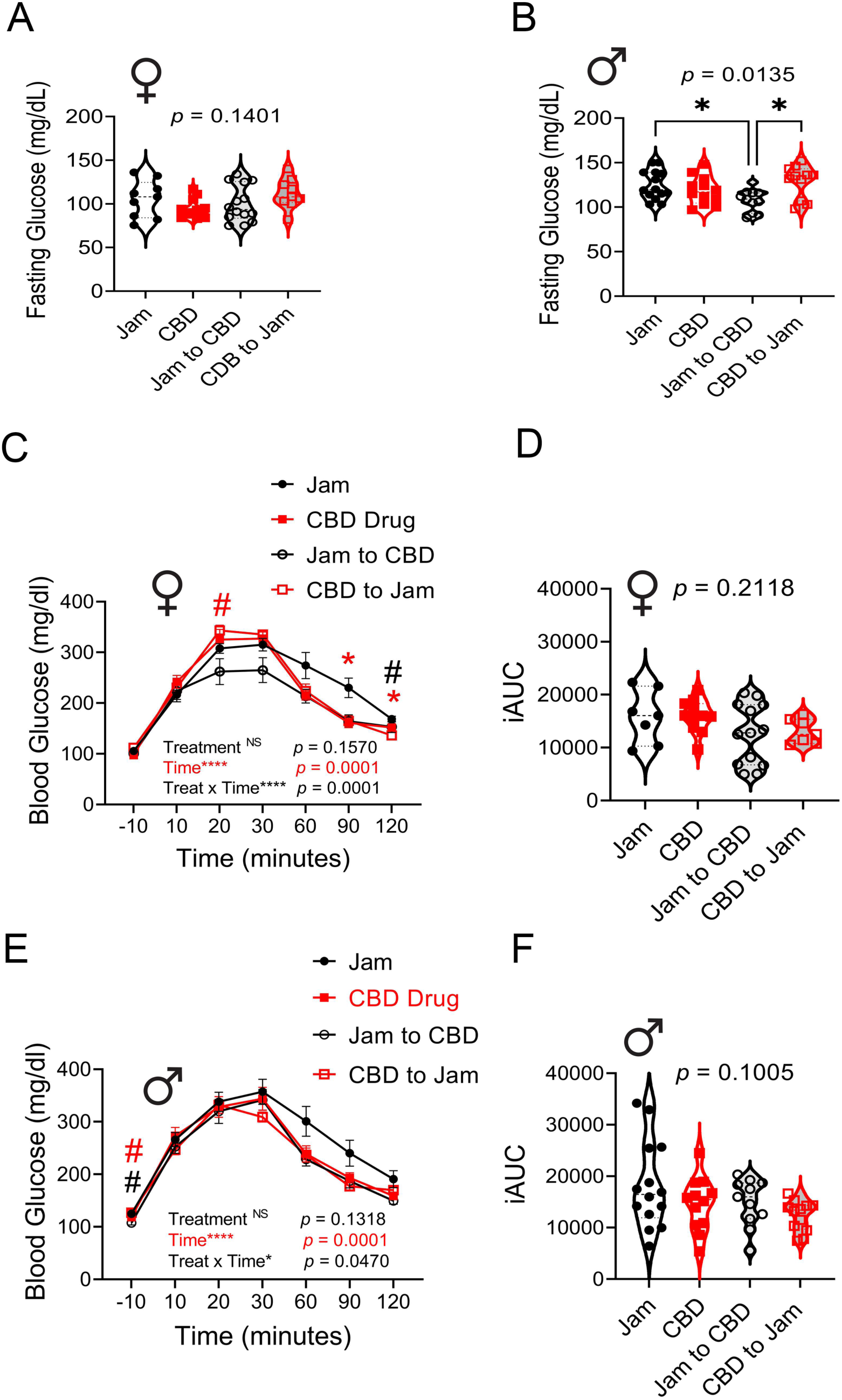
Males exposed to CBD during lactation display a lower fasting glucose and females exposed to CBD *in utero* show a faster rate of glucose clearance. Violin plot of fasting blood glucose for adult A, female and B, male offspring perinatally exposed to one of four drug treatments. Data represent individual mice with mean (± SEM), 1-way ANOVA, \**p* < 0.05. C, E, Line graph of glucose clearance and associated violet plots of D, F, integrated area under the curve (iAUC) for adult C, D, female and E, F, male offspring following an intraperitoneal glucose tolerance test (IPGTT). C, E, 2-way RM ANOVA with C, *Bonferroni* post-hoc Jam vs CBD \**p* < 0.05, #*p* < 0.05 Jam vs. CBD to Jam. E, *Bonferroni* post-hoc Jam vs. Jam to CBD #*p* < 0.05, #*p* < 0.05 Jam to CBD vs CBD to Jam. D, F, 1-way ANOVA, *p* > 0.05.

#### 3.3.3. Ingestive Behavior of Adult Offspring

Using a comprehensive laboratory animal monitoring system (CLAMS), a total of 115 adult offspring was screened for ingestive behaviors following the 4 perinatal drug treatments previously described. For clarity of presentation, complete statistical values of F and *p* are reported in **Supplementary Table 2** for both ingestive and metabolic parameters (next section). The food measures are reported in **Figure 7**, and the water measures are reported in **Figure 8**. As anticipated, mice had generally higher ingestive and metabolic activity in the dark cycle (grey shading) than during that of the light cycle (light shading), regardless of sex or drug treatment – consistent with their nocturnal behaviors (**Figures 7-9; Supplementary Table 2**). Female adult mice had no changes in meal size, number of meals, meal duration, or caloric intake in response to perinatal drug treatments in either light cycle (**Figure 7A, 7E, Supplementary Table 2**, *p* ≥ 0.05). Male adult mice that were gestationally exposed to CBD had significantly greater meal size during the dark cycle (**Figure 7B, 7F**, 1-w ANOVA, *p* = 0.0012, Bonferoni *post-hoc* test Jam vs. CBD during gestation, \*\**p* = 0.0067), and a correlate increased caloric intake (1-w ANOVA, *p* = 0.0345, Bonferoni *post-hoc* test Jam vs. CBD during gestation, \**p* = 0.0412), without changes in meal number or duration (**Figure 7C-D**, *p* ≥ 0.05). In terms of water intake, female adult mice that were exposed to CBD during lactation had significantly greater drink size during the light and dark cycles (**Figure 8A**, 1-w ANOVA, *p* = 0.0084, Bonferoni *post-hoc* test Jam vs. CBD during lactation, \**p* = 0.0126) without changes in number of drinks or even total water intake (**Figure 8C, 8E**, *p* ≥ 0.05). Male adult mice perinatally exposed to CBD had no ingestive changes with regards to water during either cycle in comparison to that of Jam treatment (**Figure 8B, 8D, 8F**, *p* ≥ 0.05). The only exception to this was a significant decrease in total water intake during the dark cycle for mice exposed to CBD during gestation, but not during lactation, compared with mice exposed to CBD perinatally (**Figure 8F**, 1-w ANOVA, *p* = 0.0358, Bonferoni *post-hoc* test CBD vs. CBD during gestation, \**p* = 0.0265).

**Fig. 7.**
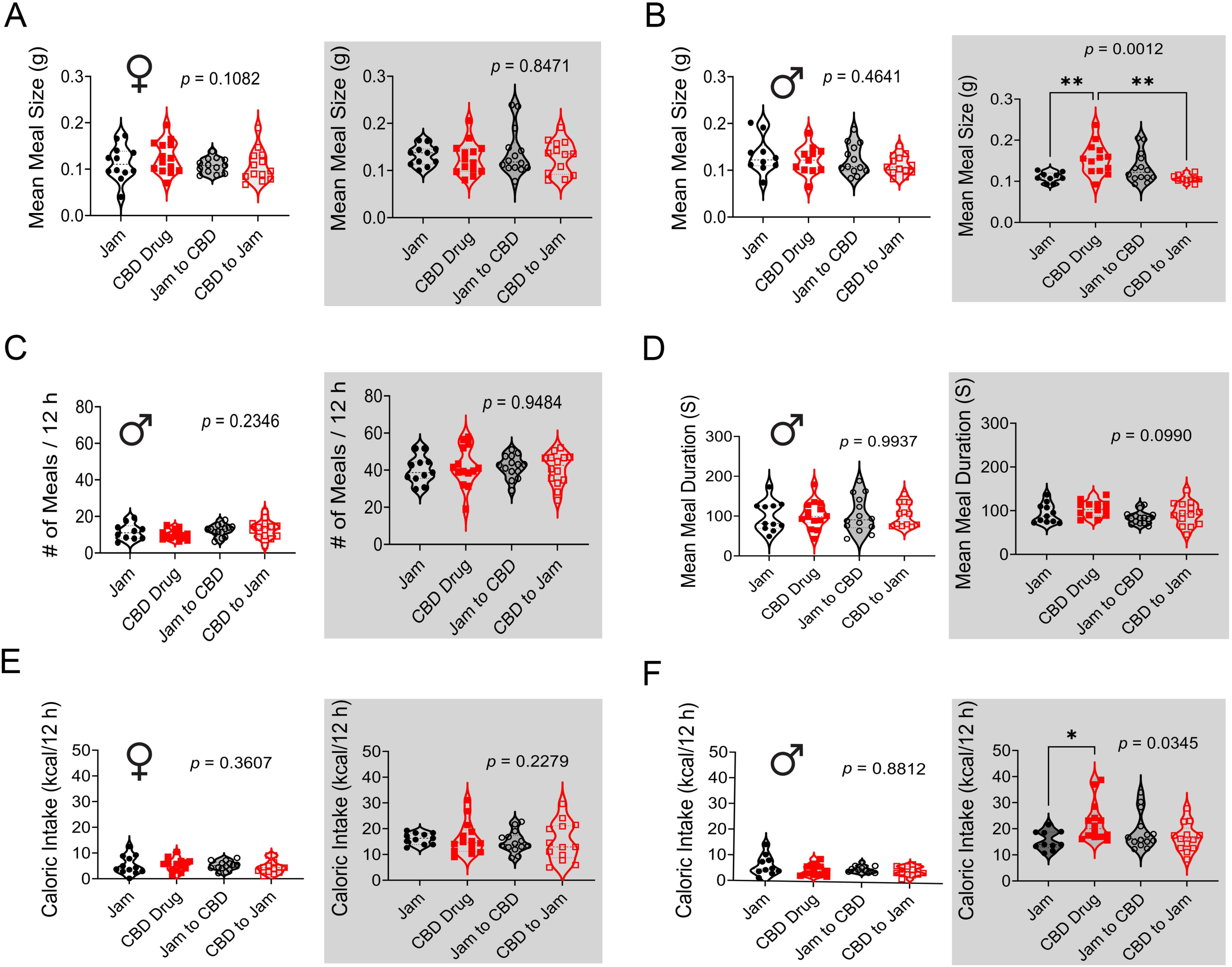
CBD exposure during gestation and lactation causes an increase in meal size and caloric intake for adult male offspring. Violin plot of A, B, meal size; C, number (#) of meals; D, meal duration; and E, F, caloric intake for female and male adult mice following perinatal exposure to one of four drug treatments. Data represent individual mice, 1-way ANOVA, *Bonferroni* post-hoc \**p* < 0.05, white background = light cycle, and gray background = dark cycle.

**Fig. 8.**
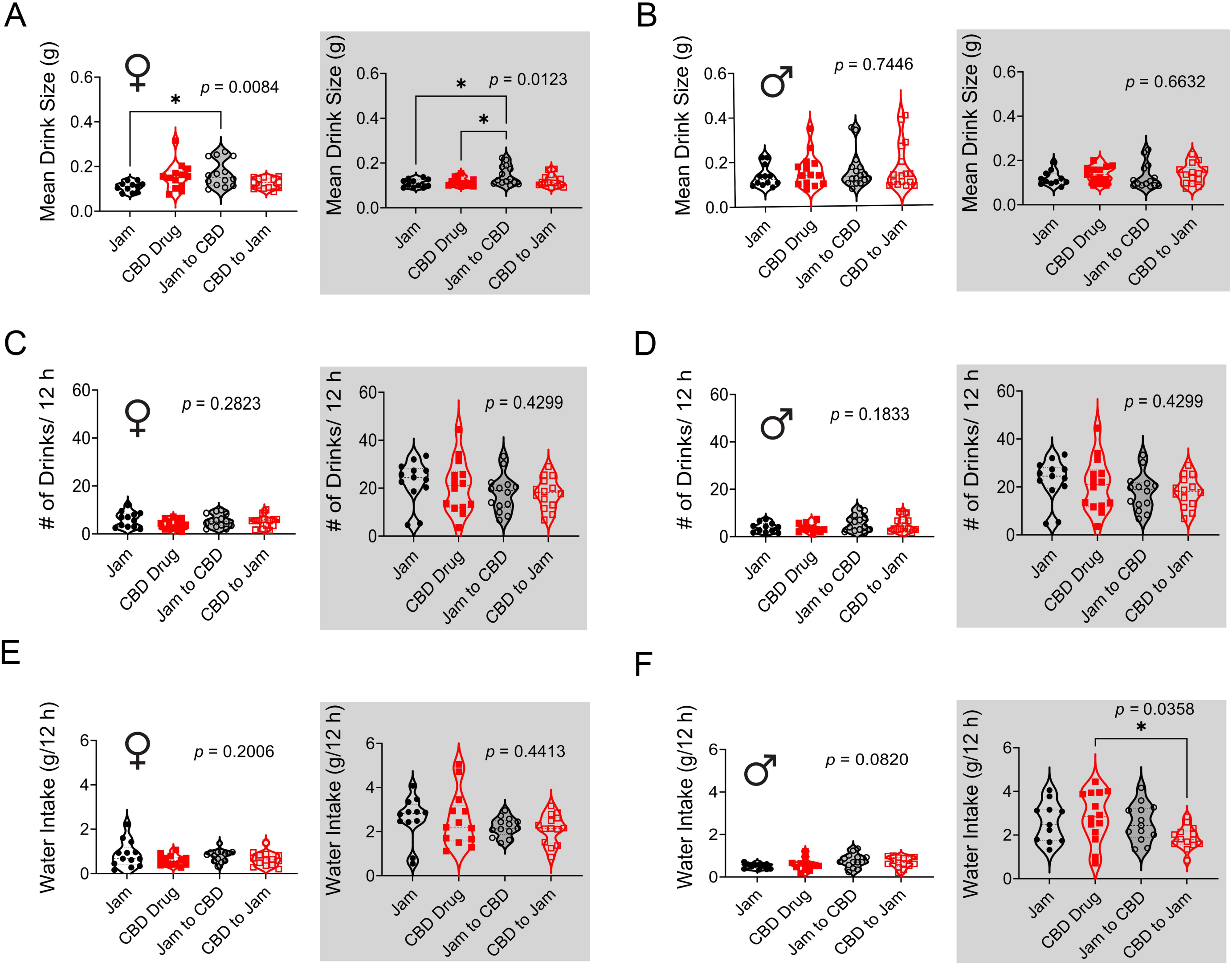
CBD exposure during lactation causes an increase in drinking size but not total water intake for adult female offspring. Violin plot of A, B, drink size; C, D, number (#) of drinks; and E, F, water intake for female (A, C, E) and male (B, D, F) adult mice following perinatal exposure to one of four drug treatments. Statistical analyses and notations as in Fig. 7.

**Fig. 9.**
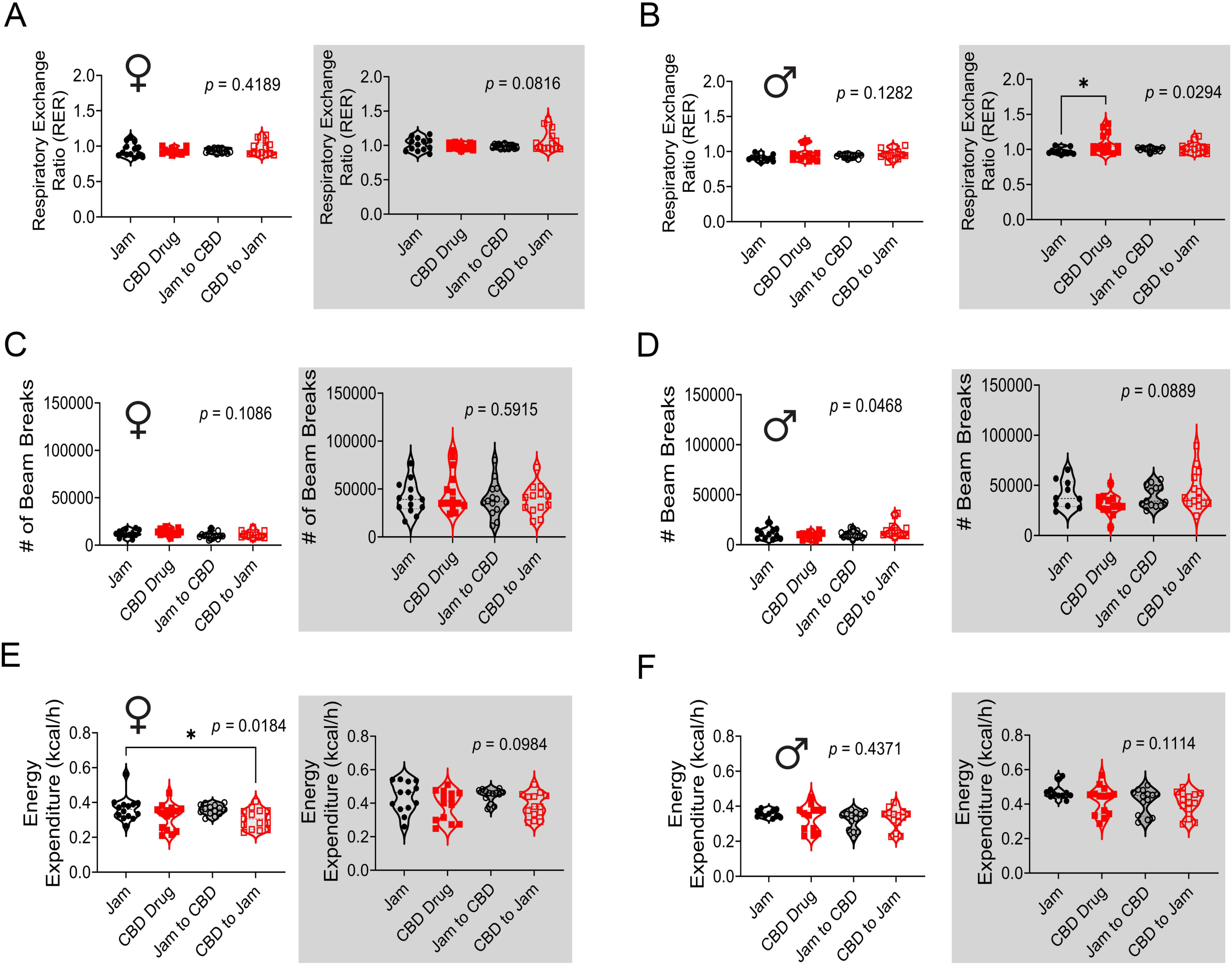
CBD exposure during gestation and lactation causes an increase in respiratory exchange ratio (RER) for adult male offspring and a slight reduction in energy expenditure (EE) for adult female offspring exposed to CBD during gestation but not lactation. Violin plot of A, B, RER; C, D, number (#) of beam breaks (ambulatory locomotion); and E, F, EE for female (A, C, E) and male (B, D, F) adult mice following perinatal exposure to one of four drug treatments. Statistical analyses and notations as in Fig. 7.

#### 3.3.4. Locomotor Activity, Energy Expenditure, and Respiratory Exchange Ratio of Adult Offspring

Also using the CLAMS, adult offspring were screened for locomotor activity, total energy expenditure (TEE), and respiratory exchange ratio (RER) following the 4 perinatal drug treatments previously described. In terms of RER, there was no main effect of drug treatment for female mice in either light cycle (**Figure 9A**, 1-w ANOVA, *p* ≥ 0.05), but there was a significant main effect of drug treatment for male mice in the dark cycle, where the ratio was greater than 1.0 for mice exposed to CBD during both gestation and lactation (**Figure 9B**, 1-w ANOVA, *p* = 0.0294; Bonferroni *post-hoc* test, Jam vs. CBD during gestation, *p* = 0.0218). In terms of locomotor activity computed as # of beam breaks during ambulatory activity in the X direction, there was no main effect of drug treatment for female mice in either light cycle (**Figure 9C**, 1-w ANOVA, *p* ≥ 0.05), but there was a modest main effect of drug treatment for male mice in the light cycle (**Figure 9D**, 1-w ANOVA, *p* = 0.0468; Bonferroni *post-hoc* test, *p* ≥ 0.05). In terms of TEE, there was a main effect of drug treatment in the light cycle for female mice (**Figure 9E**, 1-w ANOVA, *p* = 0.0184; Bonferroni *post-hoc* test, Jam vs. CBD during gestation, \**p* = 0.0429), but for male mice, there was no main effect of drug treatment in either light cycle (**Figure 9F**, 1-w ANOVA, *p* ≥ 0.05).

### 3.4 Obsessive Compulsive- and Anxiety-Like Behaviors

Following the CLAMS metabolic assessment, obsessive compulsive- and anxiety-like behaviors were measured in male and female adult offspring exposed to the previously described 4 perinatal treatments using three standard assays: the marble burying test, the light-dark box, and the elevated plus maze.

#### 3.4.1. Marble-Burying Test

Female adult mice that were exposed to CBD during lactation buried significantly more marbles (**Figure 10A**) in comparison to either Jam control or gestationally-exposed mice that were cross-fostered to a drug-free dam during lactation (**Figure 10B**, 1-w ANOVA, F (3, 24) = 4.170, *p* = 0.0164; Bonferroni *post-hoc* test, Jam vs. CBD during lactation, \**p* = 0.0157; CBD during lactation vs. CBD during gestation, \**p =*0.0477). Male adult mice did not follow this pattern observed in females, where female mice exposed gestationally to the drug, but cross-fostered to drug-free dams during lactation, buried a number of marbles that was not significantly different than Jam controls (Bonferroni *post-hoc* test, CBD to Jam vs. Jam, *p* = 0.9901). In contrast, for the male adult mice, cross-fostering to a drug-free dam during lactation did not reduce the number of marbles buried by the males compared to that of Jam control mice (**Figure 10C**, 1-w ANOVA, F (3, 30) = 5.894, *p* =0.0027; Bonferroni *post-hoc* test, Jam vs. CBD during lactation, \*\**p* = 0.0017). Therefore, there was a distinct sex-specific difference in females reversing OCD-like behavior when gestationally exposed to CBD, but provided drug free conditions during lactation, where male mice retained the OCD-like behavior even if drug free during lactation.

**Fig. 10.**
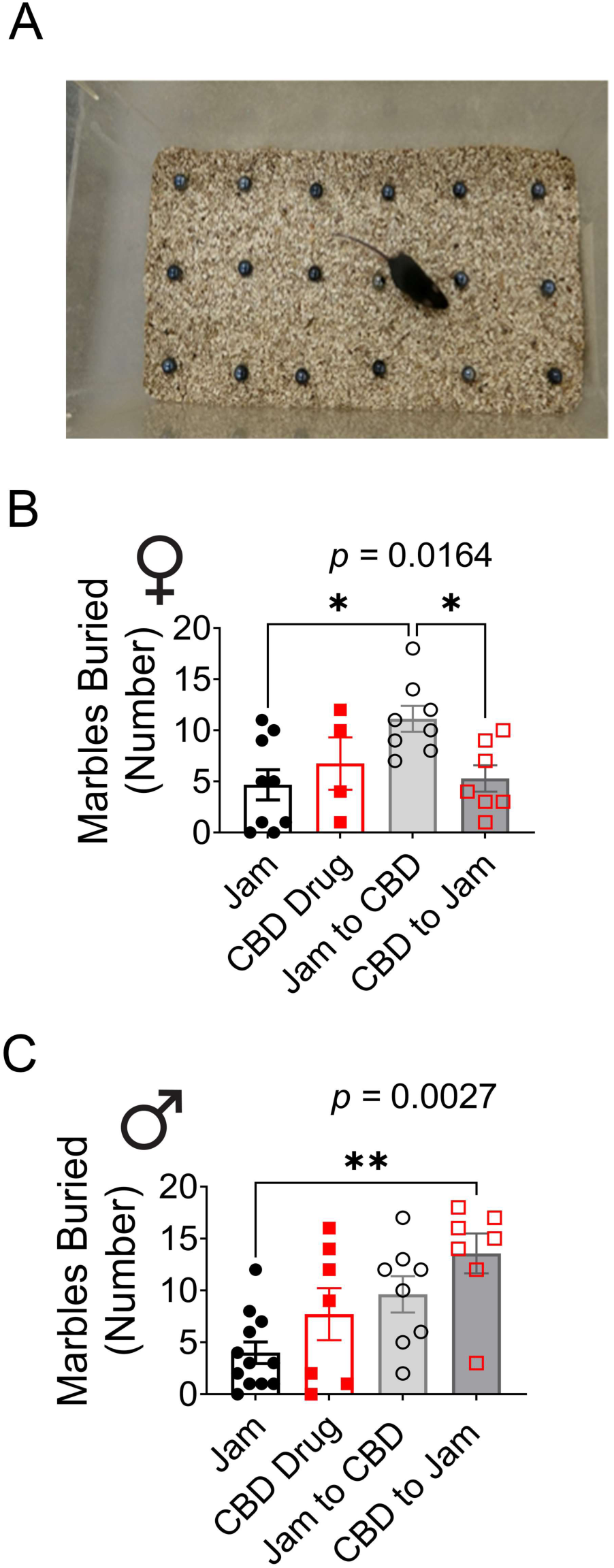
Early postnatal CBD exposure resulted in increased marble burying in adult mice, which could be reversed following cross-fostering for female, but not male, offspring. A, Photograph of the marble burying test, which serves as an indication of obsessive compulsive-like behavior. Bar graph of the number of marbles buried for B, female or C, male adult offspring following perinatal exposure to one of the four drug treatments. Data represent individual mice, 1-way ANOVA, *Bonferroni* post-hoc \**p* < 0.05, \*\**p* < 0.01.

#### 3.4.2. Light-Dark Box

Unusually, mice did not spend significantly more time in the dark compartment compared with the light compartment (**Figure 11A**), and this was true for both male and female adult offspring. This was determined using a 2-w ANOVA using compartment location as the factor (**Figure 11C, 11D**; females F(1, 48) = 0.5838, *p* = 0.5910 and males F(1, 60) = 3.753, *p* = 0.1809). This was also qualitatively observed when mouse travel tracks were plotted from mouse position determined using DeepLabCut. Because our objective was focused specifically on location x drug interactions to determine if drug treatment affected distribution of time spent in a compartment, we noted significant interactions on the LDB graphs using red font for clarity (see Interaction). The associated Bonferroni’s *post hoc p* values (*) are noted when significant for drug treatment multiple comparisons.

**Fig. 11.**
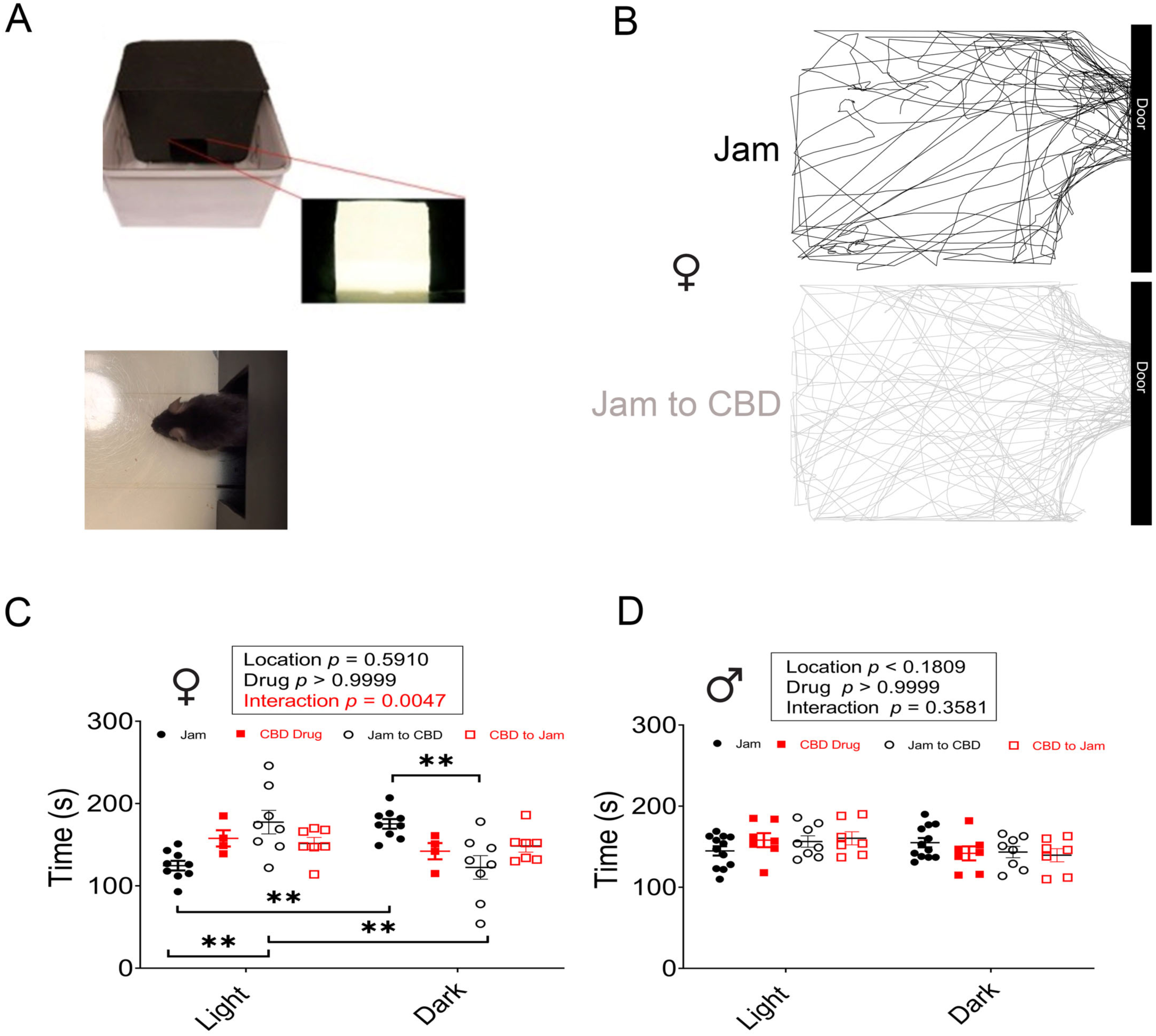
CBD exposure during lactation increases residence time in the light compartment of a light-dark box (LDB) for female adult offspring. A, Photograph of LDB apparatus with mouse positioned in the doorway. B, Representative track traces for a Jam (control) vs. Jam to CBD adult female mouse generated by DeepLabCut machine learning. Note the pattern of movement around perimeter (top panel) vs. movement through the middle of the chamber (bottom panel). Scatter plot of residence time in the two compartments for C, female and D, male adult offspring perinatally exposed to one of four drug treatments. Data represent residence time of individual mice (mean ± SEM), 2-way RM ANOVA, *Bonferroni’s* post hoc, \*\**p* < 0.01.

In female adult mice, CBD during lactation, but not during gestation, lessened anxiety-like behaviors in the LDB apparatus in comparison to the vehicle treated group (Jam). We observed a significant drug x location interaction (F (3, 48) = 11.17, *p* = 0.0047) in which female mice receiving solvent only (Jam) spent significantly more time in the dark compartment over that of the light compartment as anticipated (**Figure 11C**, Bonferroni’s *post hoc*, *p* ≤ 0.01). Adult females receiving CBD during lactation spent significantly less time in the dark compartment, and spent increased time in the light compartment, compared with those receiving vehicle (**Figure 11C**, Bonferroni’s *post hoc*, *p* ≤ 0.01). Moreover, in qualitatively observing the travel tracks generated by the DeepLabCut position data (**Figure 11B**), it was evident that the adult females that had received CBD during lactation, were traveling across the center of the open light compartment when compared to the more perimeter travel tracks observed in the Jam only mice. For the male mice, the adults not only failed to have a location preference of dark over that of the light compartment, there also was no main effect of drug treatment, nor was there a drug x location interaction (**Figure 11D**; drug as factor *p* > 0.9999, drug x location interaction *p* = 0.3581). These results suggest that early postnatal exposure of CBD (i.e. lactation), but not gestational CBD, is anxiolytic for females but ineffective for male mice.

#### 3.4.3. Elevated Plus Maze

All mice spent more time in the closed arms of the maze compared with that of the open arms or the center compartment, regardless of sex or genotype (**Figure 12A**). This was determined using a mixed RM 2-w ANOVA using arm location as the factor (all tests were highly significant *p* < 0.0001 and the individual Bonferroni *post-hoc p* values are reported on the graphs (*), see main effect – arm, **Figure 12C, 12D**). Because location was highly significant in all tests, and we wanted to focus on drug x location interactions, the significance of the latter was also denoted on the EPM graphs (#). In female adult mice, but not in male mice, there was a significant drug treatment x location interaction (**Figure 12C**, female F (6, 78) = 2.776, *p* = 0.0169; **Figure 12D**, male F (6, 96) = 1.548, *p* = 0.1708). The Bonferroni *post-hoc* test demonstrated that female mice exposed to gestational CBD (##*p* = 0.0021) or mice that were cross-fostered to drug-free dams following gestation CBD exposure (#*p* = 0.0293) spent less time in the closed arms of the maze than that of solvent-treated mice (Jam). Interestingly, representative traces generated from DeepLabCut tracking data revealed that female mice (**Figure 12B**) qualitatively showed a wider and more numerous tracks in the center and open arms for mice exposed to CBD during gestation and lactation, or during just gestation, compared to mice exposed only to vehicle (Jam). These collective results suggest that gestational CBD exposure in female mice, regardless of whether they received drug or not during lactation, is anxiolytic, whereas this effect is not observed for male mice.

**Fig. 12.**
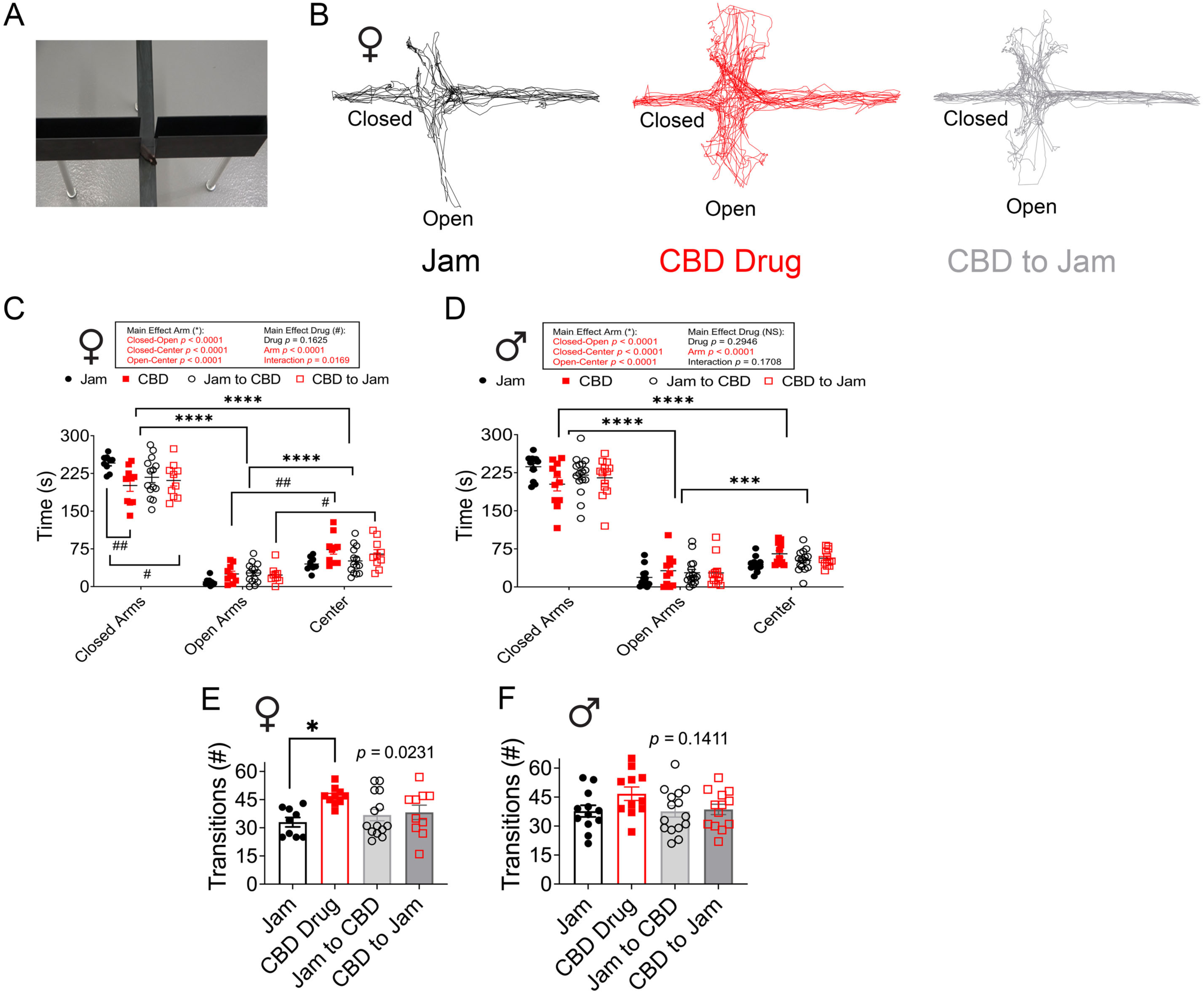
CBD exposure *in utero* decreases residence in the closed arms of the elevated plus maze (EPM) for adult female offspring and has no effect for adult male offspring. A, Photograph of the elevated plus maze (EPM) apparatus used to identify changes in heightened anxiety and avoidance of open arms (falling). B, Representative track traces for a Jam (control), CBD, vs. CBD to Jam adult female mouse generated from DeepLabCut mouse position data. Note the pattern of tight movement in control Jam (left panel), vs. that of the CBD and CBD to Jam treatments (center and right panels) where the mouse tracking has more intersections and width of movement in the center and open arms. Scatter plot of residence time in the three compartments for C, female and D, male adult offspring perinatally exposed to one of four drug treatments. Data represent residence time of individual mice (mean ± SEM), 2-way RM ANOVA, *Bonferroni’s* post hoc, using location as the factor (main effect of arm) \*\*\*\**p* < 0.0001, or using drug treatment x interaction as the factor (main effect of interaction) # < 0.05, ##*p* < 0.01. Bar graph of sum number of transitions across compartments (Transitions #) for E, female and F, male adult offspring perinatally exposed to one of four drug treatments. Data represent individual mice (mean ± SEM), 1-way ANOVA, *Bonferroni* post-hoc \**p* < 0.05.

To confirm that the CBD treatments did not cause malaise or reduction in targeted locomotor activity (as opposed to general ambulatory activity, **Figure 9C, 9D**), the total number of elevated plus maze transitions was quantified (**Figure 12E, 12F**). Rather than reduction in movement, gestational CBD exposure was found to increase the number of total transitions in female adult mice computed as the sum of all movement across all five compartments of the elevated plus maze (two open compartments, two closed compartments, and the center area) (1-way ANOVA, females F (3, 39) = 3.547; *p* = 0.0231; males F (3, 47) = 1.909; *p* = 0.1411). A Bonferroni’s post-hoc test indicated that female mice gestationally exposed to CBD had significantly more compartment transitions than that of solvent-exposed mice (\**p* = 0.0218; **Figure 12E, 12B**).

### 3.5. Object Memory and Object-based Attention Testing

#### 3.5.1. 1-Hour and 24-Hour Object Memory

Mice were required to spend greater than 20% of their observation time, but not more than 80%, with each of the two objects during the familiarization phase to ensure they had no initial bias for an object. Using this definition, 3 mice had an initial object bias during the short-term memory test and 1 mouse had an initial object bias during the long-term memory test. These mice were therefore eliminated from analysis in those respective tests. Adult mice treated perinatally with CBD demonstrated no significant change in the recognition of a novel object following familiarization with two objects (**Figure 13A**) and a subsequent wait interval of 1-hour prior to novel object presentation (short-term object memory). This was true across both sexes (**Figure 13C, 13D**, Kruskal Wallis, female *p* = 0.9211; male *p* = 0.4039). However, male adult mice that were perinatally exposed to CBD but presented with a novel object 24-hours following the familiarization with two objects, showed a change in exploration of the novel object over the familiar. The adult male mice, but not the female mice, exhibited a significantly reduced recognition index (**Figure 13E, 13F**) compared with mice receiving vehicle (Kruskal Wallis, *p* = 0.0123) and the multiple comparison test indicated the different lay between the solvent-treated mice and that mice exposed during gestation and lactation (*p* = 0.0223). Interestingly the traces generated from DeepLabCut tracking data for the male adults in the 24-hour, long-term object memory qualitatively demonstrated more circling in investigation of the novel object (cupcake) with Jam treatment vs. that of CBD treatment, where both the donut and cupcake were more equally circled and investigated (**Figure 13B**). These collective data show that CBD administration during both gestation and lactation reduced long-term memory in adult male mice.

**Fig. 13.**
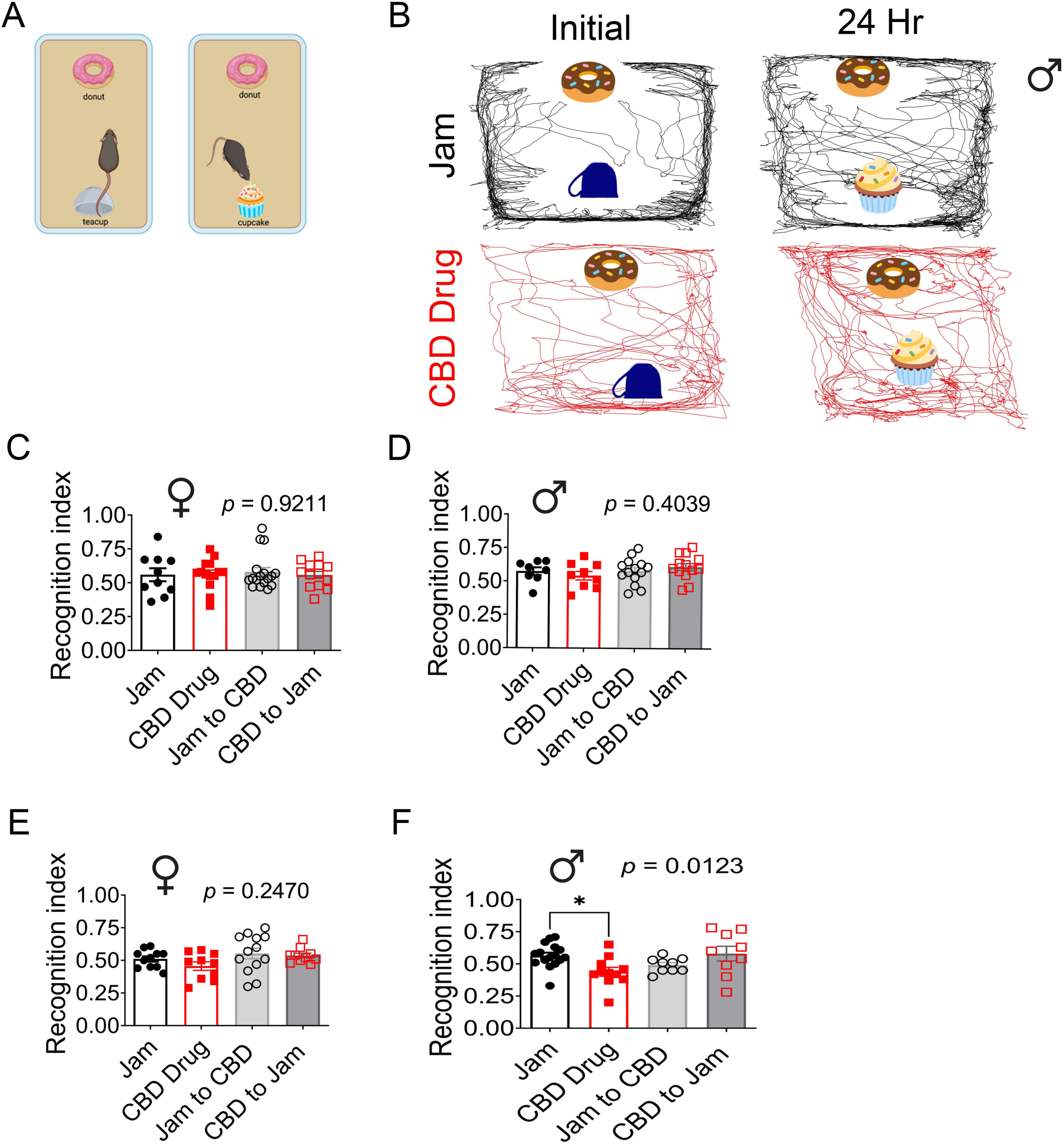
CBD exposure during gestation and lactation causes a decrease in long-term object memory in adult male offspring. A, Schematic of the object memory recognition test, where a mouse identifies the novel object (right panel; cupcake), following presentation of two initial objects (left panel) either 1 h previous (short-term object memory) or 24 h previous (long-term object memory). B, Representative track traces for a Jam (control) vs. CBD adult male mouse generated from DeepLabCut position data while performing the long-term object memory task. Note the pattern of circular movement around the identified novel object for the Jam treated mouse (cupcake, top right panel) in comparison to that of the CBD treated mouse where the track pattern is fairly equivalent between the two objects (donut and cupcake, bottom right panel). Bar graph of the recognition index (RI) of C, female and D, male adult offspring computed for 1-hr, short-term object memory vs. that computed for E, F, 24-hr, long-term object memory. Higher RI is improved memory, whereas lower RI is poorer memory. Data represent individual mice (mean ± SEM), Kruskal Wallis, \**p* < 0.05.

#### 3.5.2. Object-based Attention Test

Object-based attention tests were used to determine if gestational or early postnatal CBD exposure persistently affected the recognition of a novel object after a brief presentation of five dissimilar objects (**Figure 14A**). A lower recognition index is associated with poorer attention or ADHD-like behavior. Adult mice were observed to have no significant change in their recognition index when comparing across all 4 drug treatment groups. This was true regardless of sex of the animal (**Figure 14B, 14C – females, Figure 14D, 14E - males**; Kruskal Wallis, all *p* > 0.05).

**Fig. 14.**
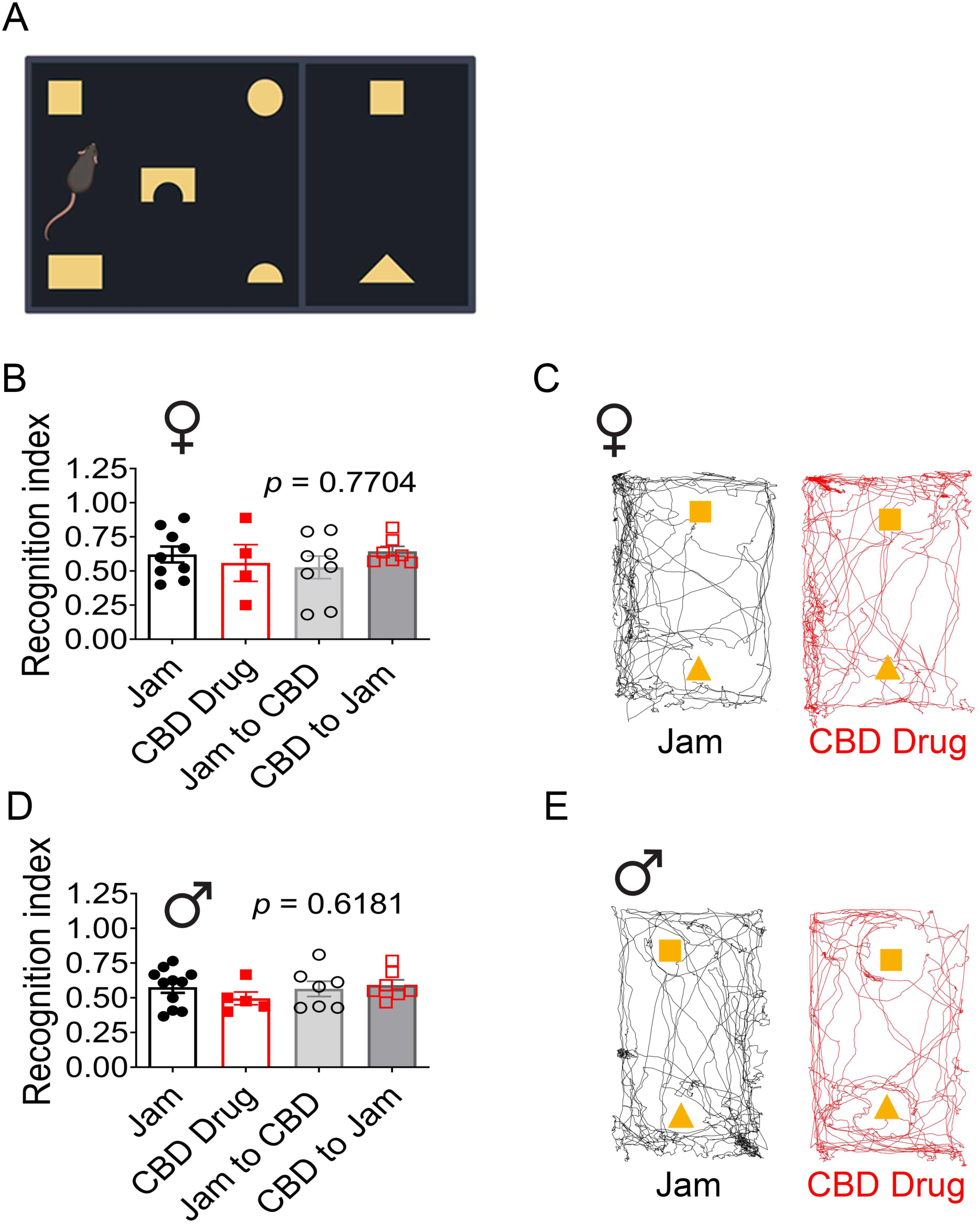
Perinatal CBD exposure. A, Schematic of an object-recognition attention test, designed to recognize attention deficit hyperactivity deficit-like activity (ADHD) in mice. Herein, a mouse is familiarized to 5 objects for a total of 5 minutes and then immediately exposed to two objects (a familiar object (square) and a novel object (triangle) for 3 minutes to determine if there is a change in attention to the novel object. B, Bar graph of the recognition index (RI) of B, female and D, male adult offspring computed for differential time spent in the second compartment with the novel object. Higher RI is improved attention, whereas lower RI is poorer attention. Data represent individual mice (mean ± SEM), Kruskal Wallis, *p* > 0.05. Representative track traces demonstrating movement in the second compartment for a Jam (control; left panel) vs. CBD (drug; right panel) C, female and E, male adult mouse generated by DeepLabCut mouse position data. Note the pattern of equivalent movement across the two objects with slight preference to novel object (triangle, < 0.50 RI) but no effect of drug treatment.

## 4. Discussion

This study developed an oral administration of CBD to pregnant dams to determine if there were persistent changes in anxiety- or attention-like behavior, or changes in object memory as adults. Because there are few studies that have determined the safety or efficacy of CBD during gestation or lactation (Wanner et al., 2021; Iezzi et al., 2022; Maciel et al., 2022; Swenson et al., 2023), our study monitored maternal health of the dams, early-life health metrics of the offspring, as well as persistent metabolic health of the offspring as adults. We discovered that oral consumption of CBD by pregnant dams did not significantly affect maternal weight gain, duration of gestation, or size of the litter, however, survival to weaning age was significantly reduced in gestationally-exposed pups that were not cross-fostered at birth to drug-free dams.

Perinatal exposure to CBD caused a change in early postnatal developmental body weight that was sex dependent, and early postnatal CBD exposure during lactation lowered fasting glucose levels in male adult mice. Gestational exposure to CBD caused male mice to increase meal size and caloric intake without significant disruption to the number or duration of meals, whereas early postnatal exposure to CBD during lactation enhanced drinking size in female mice without significant disruption to total water intake. The greatest metabolic modification attributed to perinatal CBD exposure appears to be a change in respiratory exchange ratio (RER), whereby there was an increase in carbohydrate use as fuel in male mice, specifically in the dark cycle. In terms of adult behaviors, early postnatal exposure to CBD during lactation evoked an increase in OCD-like behavior in female mice that could be mitigated by cross-fostering to a drug-free dam, but not for male mice, who retained an OCD-like behavior as adults following gestational exposure. Two different anxiety tasks support the observation that CBD perinatal exposure is anxiolytic for female, but not for male, mice. Finally, adult mice are not affected by CBD perinatal exposure in terms of ADHD-like behavior or short-term object memory, however, gestational CBD exposure for male mice was found to decrease long-term object memory.

Our study design advantageously utilized cross-fostering to separate the perinatal effects of CBD during gestation, during lactation, or both. Most preclinical studies have stopped cannabinoid administration at, or just prior, to birth, which in the rodent brain, roughly corresponds developmentally to early third trimester in human brain development (Pressler and Auvin, 2013). Metrics such as reduced fasting glucose levels, increased mean drink size, changes in OCD-like behavior, and LDB-tested lessoned anxiety-like behavior were uncovered during exposure to the drug during lactation specifically, which may not have been discovered with only *in utero* administration. These types of drug-induced modifications in metabolism or behavior in the animal as adults, may indicate a late trimester or early developmental critical window when there is more sensitivity to CBD, and showed sexually dimorphic differences.

We believe our study may be the first to use voluntary CBD drug administration as opposed to oral gavage (Wanner et al., 2021; Swenson et al., 2023). Given the fact that nearly all our monitored dams (56 out of 57) successfully acclimated to eating jam from the dish, and within one week, consumed CBD/jam within 15 seconds, indicates that this route of administration not only mimicked consumption of edibles in humans, but also allowed synchronization across animal cohorts for uniform timing of the dose, prior to examining for a subsequent task, tissue collection, or behavior. Moreover, this form of oral consumption is very amendable for testing during pregnancy because it minimizes stress. Other routes of administration such as intraperitoneal injection, subcutaneous injection, and oral gavage (Iezzi et al., 2022; Wanner et al., 2021; Maciel et al., 2022; Swenson et al., 2023) can be associated with decreased litter size and increased uterine reabsorption events when used during pregnancy (Black et al., 2023). The downside of an oral administrative route is its slower absorption; presenting bioavailability ranging from 4 to 12%, which is dictated by gastrointestinal degradation and extensive first-pass metabolism (Grotenhermen, 2003; McGilveray, 2005; Xu et al., 2019). Given the poorer bioavailability of CBD across the digestive tract compared with that of intravenous, IP, or SQ routes of administration (Grotenhermen, 2003; McGilveray, 2005; Xu et al., 2019), our results using oral administration of 100 mg/kg CBD in strawberry jam, showed a lower peak concentration but a similar kinetic rise (422 ng/ml) within 30 minutes that fell rapidly to below our detection levels (10 ng/ml) within 4 hours. This is consistent with approximately 14% dose administration by IP injection and comparable with other studies that evaluated several routes of CBD administration (Deiana et al., 2012). Soni et al., 2023 report that the use of an aqueous vehicle vs. that of sesame seed oil by IP injection effects the speed at which plasma concentrations of CBD rise and fall (aqueous injections result in high peaks that rapidly decrease) (Soni et al., 2023). In future studies we should determine if the combination of the drug with fruit/sucrose vehicle (strawberry jam) versus oil alters the kinetic pattern of drug concentration in the plasma. Ochiai et al., 2021 report rates of CBD transfer to fetal brain, liver, and gastrointestinal tract within 15 minutes of dam injection, which is consistent with the findings that typical oxidation of CBD by the liver using CYP3A is poor in the fetus that has very low concentrations of this metabolizing enzyme (Ochiai et al., 2021).

It is difficult to judge the physiological dose of CBD administration for a perinatal study because routes of administration affect pharmacokinetics and influence bioavailability; cannabinoids can be transferred through the BBB, through breast milk, and are differentially transferred across organs; plus, one must consider metabolic factors and size of the animal model (Reagan-Shaw et al., 2008). Our study used a dose of 100 mg/kg in mice, which can be compared with the only FDA-approved use of CBD – a dose of 25 mg/kg per day for epilepsy in humans (Silvestro et al., 2019; Abu-Sawwa and Stehling, 2020). This converts to 175 mg/kg in mice when considering metabolic and size differentials between humans and mice (Reagan-Shaw et al., 2008). Therefore, we are well below this clinical dose in our study, and of course we are administering to dams which transfer drug to the offspring. Although our study did not measure CBD levels in offspring, the closest study and route of administration similar to our own is that of Swenson et al., 2023, who found that 50 mg/kg CBD prepared in sunflower oil and delivered by oral gavage, crosses the placenta and is found in pup plasma through postnatal day 8 if administration was stopped at birth (Wanner et al., 2021; Swenson et al., 2023). Albeit using a subcutaneous CBD route of administration, another study using a lower 3 mg/kg dose report significant levels of CBD in embryonic brain, and that levels in males were significantly less than in females, which may explain some of the sex-dependent behavioral differences we noticed in our study (Maciel et al., 2022).

Of interest to our study that used a cross-fostering design and observed significant effects of CBD attributed to exposure during lactation, are studies that measure bioavailability of CBD in breast milk. Soni et al 2023 measured CBD transmitted to pups, where 50 mg/kg CBD (80% tween/PBS) was administered by IP to a lactating dam, which was detected in pup plasma 18 h post injection at a level ranging between 10-15 ng/ml (Soni et al., 2023). When examining the breastmilk of CBD-treated dams, breast milk contained a significant reduction in N-acyl methionine and free fatty acids, which have been shown to promote litter weight gain and protein synthesis (Liu et al., 2017; Johnson et al., 2022). A reduction of these weight promoting macromolecules in breastmilk could be a contributing factor for the lower body weights we observed in the early postnatal male mice (P10-P21), especially prominent in pups that had CBD only administered during lactation, and which began to better weight stabilize following their transition to solid food (following weaning). In all, few pharmacokinetic studies of CBD during the perinatal period exist yet are necessary to understand relevant exposure consequences during early embryonic and postnatal development.

We observed a significant decrease in survival to weaning age (P23) for offspring that were exposed to CBD during gestation and lactation. While we do not know the mechanism of the poorer survival following exposure, there are two variables that should be considered – 1) the use of synthetic vs. plant-derived CBD, and 2) dose and duration of exposure. Our study employed the use of synthetic CBD to separate the effect of CBD from other cannabinoids present in a plant extract or inconsistencies between batch extractions. Some concern has been raised as to the safety and efficacy of synthetic cannabinoids (Cohen and Weinstein, 2018) that have been attributed to higher binding affinities to endocannabinoid targets. Mass spectrometry testing from the National Institute on Drug Abuse, however, confirms that synthetic and plant-derived CBD contain the same molecule (Insys Therapeutics, 2014). Research comparing efficacy in the treatment of epilepsy and inflammation has also shown that plant-derived and synthetic CBD are equivalent (Klotz et al., 2019; Maguire et al., 2021). In terms of duration of perinatal exposure, Swenson et al., 2023, found no effect on pup survival following 50 mg/kg of fetal CBD exposure administered by oral gavage (Swenson et al., 2023), however, the duration of perinatal exposure was from embryonic days 5 to 18, and exposure during our study was a total of 58 days. The only other study that reported pup survival following perinatal CBD exposure was in rats, as opposed to mice, also with a lower concentration (3 mg/kg in rats corresponding to a 22 mg/kg dose for mice; Vanin et al., 2023) and shorter exposure (embryonic 6 to birth), and with monitoring only taking place until postnatal day 4. Shorter exposure time or lower doses may have better neonatal outcomes suggesting a possible dose toxicity, and indicating future dose-responsivity studies are warranted to determine if there is a safe dosage for CBD use during pregnancy.

Our data show that litter size and maternal weight gain do not appear to be altered following perinatal exposure to CBD, similar to that reported by Swenson et al., 2023. We found that initial P1 body weight was equivalent across treatments, suggesting lack of drug-induced developmental changes that would affect gross body growth *in utero*. However, we found that exposure to CBD *in utero* or during lactation appeared to be detrimental to early postnatal thriving, especially for male offspring. Male offspring that experienced CBD exposure during gestation and lactation or only during lactation, had significantly reduced body weight compared to that of vehicle control offspring, and this persisted through early postnatal development - P10 to P21. This observed failure to thrive during early postnatal stages may be linked to drug-induced changes in milk composition (Johnson et al., 2022) as previously mentioned and does not appear to be attributed to poor or altered maternal behaviors. This is an important distinction because poor maternal care has been linked to neuropsychiatric disease in humans and neurodevelopmental problems in rodents (Shin et al., 2013; Bridges, 2015; Sacks et al., 2017). Iezzi et al., 2022 reported sex-specific changes in pup vocalizations following CBD administration during a window of gestational time (embryonic days 5 to 18), but they found no resultant changes in maternal pup retrieval. We also did not observe any significant change in maternal pup retrieval by several metrics (first latency and full retrieval), nor did we qualitatively observe poor maternal behaviors (i.e. failure to prepare a nest, lack of milk ribbons, walking on pups, or scattering pups outside the nest). There is strong evidence through a recent PubMed meta-analysis (years 2014-2023) performed by Brianna Moore (2024) that perinatal exposure to Δ9-THC in human and in animal studies results in low body weight (LBW) and an increase in NICU admission with an acceleration of weight or “catch up” in early development that could result in poorer metabolic health (Moore, 2024). There are some studies that report males being more susceptible to this gestational drug exposure (Gunn et al., 2016; Benevenuto et al., 2017). This is very strikingly similar to what we observed in the male offspring, that appeared to better stabilize their weights by P60. Ignatowska-Jankowska et al. found that a daily, two-week CBD administration via IP injection produces a significant decrease in body weight in adult male rats (Ignatowska-Jankowska et al., 2011). Of the 4 published works that report perinatal CBD exposure on health outcomes (Wanner et al., 2021; Iezzi et al., 2022; Maciel et al., 2022; Swenson et al., 2023), only one measured early postnatal pup body weight (Iezzi et al., 2022), and while they found it was also sex-dependent – affecting males – they report as much as 1-2 g heavier body weights from ages P10 to P22. A complication in interpretation again lies in the differential experimental variables – namely, route of administration, total duration of drug treatment, and the time since last CBD administration. In our study, we administered drug 2 weeks prior to mating and then throughout gestation and lactation (a total of 58 days). Our reported reduction in male body weight (until P21) was for mice that were still receiving CBD, whereas those in the Iezzi et al., 2022 study had been drug free several days before birth (E5 to E18 was drug interval) and all during the postnatal development period. In our study, if gestationally exposed pups were cross-fostered to drug-free dams during lactation (CBD to Jam), their body weight was not significantly different than Jam vehicle. Moreover, following weaning, when male pups were completely drug-free for 40 days, their body weights largely stabilized; typical of a rebound-like activity (Nashed et al., 2020). We observed that the only remaining treatment resulting in reduced body weight in the males as full adults (P60), was for pups that were exposed to CBD throughout lactation. It is possible that the postnatal period represents a critical developmental period during which weight rebound is not achievable to normal levels.

It is well known that a reduction in body weight is correlated to improved fasting glucose levels in both diabetic and non-diabetic individuals, due primarily to an increased sensitivity to insulin (Kong et al., 2020). This is consistent with our observed changes for male mice, which exhibited a decrease in postnatal body weight and a measured reduction in fasting glucose levels following treatment with CBD during gestation or gestation and lactation. This is counter to what was reported in gestationally-exposed male rats (Vanin et al., 2023), for which IP injection of CBD (E6 to birth) caused long-term liver damage, glucose intolerance, and no change in fasting glucose levels. In our study, while there was no change in overall glucose clearance for either sex (iAUC data), females offspring demonstrated a faster kinetics of clearing a glucose challenge. Interestingly, in human studies, there was no effect of prenatal cannabis on fasting glucose or insulin in either male or female offspring (Moore et al., 2022).

Adult male offspring exposed to CBD during gestation and lactation had an increased meal size and increased caloric intake. This is opposite of what we would have expected. Given the role of CB1 and CB2 receptors in feeding behavior (Tarragon and Moreno, 2019), and CBD’s known activity as an antagonist of these receptors, we would anticipate that administration of CBD would decrease food intake (Scopinho et al., 2011; Farrimond et al., 2012; Bielawiec et al., 2020). Perhaps CBD functions developmentally in a compensatory mechanism to regain body weight from the low body weight of the early postnatal stages (P10-P21). This may suggest that when gestationally administered, CBD continues to function as an anti-obesity agent (Ignatowska-Jankowska et al., 2011; Farrimond et al., 2012; Tarragon and Moreno, 2019; Bielawiec et al., 2020). Interestingly, in human studies, fetal cannabis exposure leads to higher fat mass, fat-free mass, and adiposity (Moore et al., 2022). It is also interesting to note the concomitant elevation in RER in the dark cycle that is reflective of a greater use of carbohydrates over that of fat as fuel in the adult males exposed to CBD during gestation and lactation. A greater use of carbohydrates as fuel would be advantageous for offspring with a lowered body weight since it would help them gain weight more quickly. A RER value greater than 1.0 can not only indicate exclusive use of carbohydrates as fuel but can alternatively indicate fatigue following enhanced exercise and oxygen debt (Katch et al., 2011). This does not appear to be a likely interpretation in our study because we did not observe either an increased ambulatory activity or increased locomotion with this drug treatment. In humans, a single CBD exposure fails to change RER following a meal but does increase in RER following exercise (Abbotts et al., 2022; Sahinovic et al., 2022). Previous studies examining exposure in adult mice found that long-term (or chronic) CBD treatment can decrease ambulatory activity, while short-term (or single exposure) treatment had no effect on locomotion (Schleicher et al., 2019; Viudez-Martínez et al., 2019; Kaplan et al., 2021; Calapai et al., 2022). In our study, mice were chronically exposed to CBD for up to 58 days, and metabolically profiled following approximately 60 days of drug free development. This suggests chronic perinatal CBD effects on locomotor activity could disappear after drug administration is stopped.

Our study indicated that adult females exposed to CBD during lactation or early postnatal development had a persistent increase in drinking size independent of light cycle. While there has been evidence that THC reduces salivation by interacting with the CB1 receptor, thereby reducing acetylcholine production, and downregulating basal salivation (Andreis et al., 2022) – and this effect is sex independent - our increased drinking was only observed in females. Because CBD is an antagonist, rather than an agonist for CB1 receptor, such a receptor interaction would be predictive of an upregulation of salivation, rather than thirst producing. Moreover, an increased drinking size did not lead to an overall change in daily water consumption. Perinatal CBD exposure may have developmentally altered drinking behavior (size) in a sex-dependent fashion and may be distinct from single CBD administration (Sofia and Knobloch, 1976), where water intake is decreased.

In terms of adult behaviors, it is interesting that early exposure to CBD increased obsessive compulsive-like behaviors (marble burying) in both males and females, which could be reversed in females if cross-fostered to a drug-free dam, but not reversed for males. This suggests a sexual dimorphic developmental response to CBD; males exposed to CBD *in utero* retained this behavioral trait as adults even when the drug-free period was initiated earlier and with control maternal nurturing. It is interesting that obsessive compulsive behaviors in humans are thought to present earlier in childhood for males but are more common in females in adolescence and adulthood (Mathes et al., 2019).We and others (Nardo et al., 2014; Murphy et al., 2017; Huffstetler et al., 2023) have reported anti-obsessive compulsive behavior in male mice following either acute or chronic administration of CBD in adults. Deiana et al., 2012, reported an increase in obsessive compulsive-like behavior in mice linked to the pharmacokinetics of CBD - after 6 hours of progressive drug elimination in plasma, normal marble-burying behaviors were observed, indicating behavioral outcomes may also be determined by a threshold. It is well known that sensitivity to CBD is greater in fetal stages over that of adults (Mulligan and Hamre, 2023; Swenson et al., 2023). In comparison with these collective findings in adults, exposure to CBD developmentally may lead to more long-lasting changes (perhaps changes in gene expression, Wanner et al., 2021), and ones that are sexually dimorphic, affecting males to a greater extent than females.

In two anxiety tests, the light-dark box (LDB) and elevated plus maze (EPM), perinatal exposure to CBD functioned as an anxiolytic in female, but not male mice. In the female mice, for the LDB tests, the greatest effect in lowering anxiety-like behavior was for drug exposure during lactation, whereas for the EPM, both exposure during gestation or throughout gestation and lactation had a significant ability to lower anxiety. Swenson et al., 2023, who orally gavaged pregnant dams to expose offspring from E5 to E18, did not observe any changes in anxiety-like behavior in adults for either sex in the LDB or the elevated zero maze. Because our study made use of cross-fostering, we may have been able to observe anxiety-like behaviors attributed to the sensitive period of lactation (early postnatal development) for which they did not test or examine, and again, this behavioral change was sex dependent, this time for females. CBD is known to bind and activate endocannabinoid receptors expressed in the fetal brain that are important for brain development, including serotonin receptors (5HT1a), and excessive activation of these receptors can disrupt neurodevelopment. Particularly, overexpression of 5HT1a during mouse fetal and early postnatal development has been shown to decrease anxiety-like behaviors (Bert et al., 2008). Future studies could examine excessive or sexually-differential activation of these pathways in the fetal brain exposed perinatally to oral CBD administration. The male mice exposed to jam (control solvent) exhibited an effect of location (sidedness or arm) for the EPM (preferred closed arms) but did not reach significance (*p* = 0.1809) in the LDB (no preference for dark over light) – which was unexplained. Because we have run large sample sizes of wildtype and transgenic mice in LDB studies (Marks et al., 2009; Huang et al., 2018b; Kolling et al., 2022; Huffstetler et al., 2023) and have never observed this in control animals, we can only conclude that administration of jam must cause some type of flattening of the expected basal location effect in the male offspring.

Increased levels of anxiety or the presence of generalized anxiety disorder is known to decrease memory or cognition (Lukasik et al., 2019). Despite males having no change in anxiety-like behavior as adults, we did observe a decrease in long-term object memory. We have previously demonstrated that acute, single administration of CBD in adults causes a similar decrease in long-term object memory in male mice (Huffstetler et al., 2023). Interestingly, CBD binds and activates the potassium channel, Kv7.2/3, which is expressed in the brain throughout embryonic and postnatal development (Dirkx et al., 2020). Alterations of fetal Kv7.2/3 activity has been known to cause cognitive and memory deficits in mice (Zhou et al., 2011), and thus may be a future target to explore as a potential mechanism for causing alternation in long-term object memory following perinatal CBD. Fewer studies have examined cognitive or attention tasks following CBD administration (Huffstetler et al., 2023) and none have examined this behavior following CBD perinatal exposure. Similar to our findings that perinatal CBD exposure had no effects on ADHD-like behavior in adult offspring, acute, single exposure of CBD in rodents failed to change attention or ADHD-like behaviors (Huffstetler et al., 2023; Moore et al., 2023). The lack of effect of CBD contrasts with what is well-studied in rodents with regards to that of THC and impaired attention (Moore et al., 2023; Sarikahya et al., 2023), yet larger meta-analyses in humans have reported cognitive changes that fall within normal ranges when adjusted for age and education (Torres et al., 2020).

Our study has not yet examined a mechanism for the long-lasting metabolic and behavioral changes we measured in adults that were exposed to CBD *in utero.* A possible cause of such long-lasting changes could be either changes in gene expression or altered signaling pathways. Wanner et al. has shown that perinatal CBD exposure causes changes in patterns of DNA methylation (Wanner et al., 2021), suggesting epigenetic modifications during fetal development with drug exposure. Epigenetic modifications have also been implicated in metabolic reprogramming and transcriptional regulation of metabolic enzymes, which may explain some of the metabolic changes we noticed (Huo et al., 2021). Sarikahya et al. found changes in protein expression concomitant with an increase in marble-burying following prenatal cannabis exposure (Sarikahya et al., 2023). Additional studies are needed to examine molecular changes in gene expression and neural activity following CBD exposure to elucidate mechanisms leading to behavioral and metabolic changes.

## 6. Conclusions

In conclusion, through trained, voluntary oral administration of CBD to pregnant dams, we found that CBD caused persistent changes in metabolism and behavior when raised to an adult. CBD exposure did not significantly alter maternal weight gain, gestation length, or litter size. However, it troublingly decreased the survival of offspring to weaning, reduced the weight of male offspring, increased their use of carbohydrates as fuel, and lowered their fasting glucose levels. Our study separates important periods of development and investigates the influence of CBD exposure during gestation, lactation, or both, which has previously not been examined. Simultaneously we examined long-term effects in male and female offspring as adults to discover sex-dependent changes in behavior and metabolism. CBD exposure during what may be a developmental critical period during lactation, and analogous to third trimester in humans, resulted in reduction of fasting glucose levels and increased meal size for male mice, while females, it resulted in an increase in drink size. Female mice exhibited increased OCD-like behavior and decreased anxiety-like behavior if exposed to CBD during lactation, whereas male mice exhibited OCD-like behavior to a significant extent when exposed gestationally, which was not mitigated by cross-fostering to a drug-free dam and had no change in anxiety attributed to CBD perinatal exposure. Perinatal exposure to CBD did not affect attention tasks, but selectively decreased long-term object memory in males raised to adults. Our study highlights the need for further research on the developmental impacts of CBD, especially considering the growing use of CBD during pregnancy, and in establishing safe doses of CBD during pregnancy and tighter regulation of CBD products given the potential negative outcomes for offspring in this preclinical study.

## DATA AVAILABILITY

All video recordings acquired during our behavioral experiments can be made available through contact of the communicating author. Sharing of custom scripts for DeepLabCut and representative traces are also freely available through contact of the communicating author.

## ACKNOWLEDGEMENTS

We would like to thank Drs. Kathleen Harper, Jaime White-James, and William Hill for their care and oversight of the FSU vivarium and their extra assistance in the logistics of our behavioral phenotyping. We would like to thank Mr. Ken Snoke and Bob Schmidt of Emerald Scientific for their time and knowledge that they relayed concerning cannabidiol purity, synthesis, handling, and applications. This work was supported by the Consortium for Medical Marijuana Clinical Outcomes Research (MMCOR) of the State of Florida, the Higher Education Emergency Relief Funding (HEERF) through the CARES act, the FSU Research & Creativity Council through a Planning Grant, the Bess Ward Scholarship at Florida State University Honors Program. The Mass Spectrometry Research and Education Center at the University of Florida (NIH S10 OD021758) supported our CBD determination in the maternal plasma.

## ABBREVIATIONS

1-w ANOVA: one-way analysis of variance
2-w ANOVA: two-way analysis of variance
ADHD: attention deficit hyperactivity disorder
AM-251: inverse agonist at the CB_1_ cannabinoid receptor
ARRIVE: Animal Research: Reporting of *In Vivo* Experiments
AVMA: American Veterinary Medicine Association
BW: body weight
CBD: cannabidiol
CB1 (CB_1_-R): cannabinoid-1 receptor
CLAMS^TM^: comprehensive laboratory animal monitoring system
cm: centimeters
CNS: central nervous system
dF: degrees of freedom
EDTA: ethylenediaminetetraacetic acid
EE: energy expenditure
EPM: elevated plus maze
ESI: electrospray ionization
EtOH: ethanol
F statistic: F-value, ratio of two variances
F_max_ Test: Hartley’s test, Hartley’s F_max_, tests for homogeneity of variance
FO: familiar object
g: grams
GAD: generalized anxiety disorder
GMP: good manufacturing practices
H: height
h/hr: hour
IACUC: Institutional Animal Care and Use Committee
iAUC: incremental area under the curve
IP: intraperitoneal
IPGTT: intraperitoneal glucose tolerance test
kcal: kilocalories
kg: kilograms
kV: kilovolt
LDB: light-dark box
M: molar
mCPP: meta-chloro-phenyl-piperazine, serotonin agonist
mg: milligrams
min: minutes
ml: milliliters
mm: millimeters
mM: millimolar
m/z range: mass-to-charge ratio
n: number
ng: nanograms
NIH: National Institute of Health
NO: novel object
OBS: object bias score
OCD: obsessive compulsive disorder
OFT: open field test
P: postnatal day
RER: respiratory exchange ratio
RI: recognition index
RM: repeated measure two-way analysis of variance
RPM: revolutions per minute
s: seconds
TEE: total energy expenditure
THC: Δ9-tetrahydrocannabinol
TV: television
μl: microliters
μm: micrometers
VCO_2_: volume carbon dioxide produced
VO_2_: volume oxygen consumed
WT: wildtype
YFP: Thy1-YFP, mice with green labeled olfactory bulb neurons, mitral YFP

